# Systems metabolic engineering of *Corynebacterium glutamicum* for efficient L-tryptophan production

**DOI:** 10.1101/2024.11.04.621991

**Authors:** Yufei Dong, Zhen Chen

## Abstract

Corynebacterium glutamicum is a versatile industrial microorganism for producing various amino acids. However, there have been no reports of well-defined C. glutamicum strains capable of hyperproducing L-tryptophan. This study presents a comprehensive metabolic engineering approach to establish robust C. glutamicum strains for L-tryptophan biosynthesis, including: (1) identification of potential targets by enzyme-constrained genome-scale modeling; (2) enhancement of the L-tryptophan biosynthetic pathway; (3) reconfiguration of central metabolic pathways; (4) identification of metabolic bottlenecks through comparative metabolome analysis; (5) engineering of the transport system, shikimate pathway, and precursor supply; and (6) repression of competing pathways and iterative optimization of key targets. The resulting C. glutamicum strain achieved a remarkable L-tryptophan titer of 50.5 g/L in 48h with a yield of 0.17 g/g glucose in fed-batch fermentation. This study highlights the efficacy of integrating computational modeling with systems metabolic engineering for significantly enhancing the production capabilities of industrial microorganisms.

## 1. Introduction

L-tryptophan is an essential amino acid for both animals and human, which plays vital roles in protein synthesis and metabolic regulation. L-tryptophan and its derivatives [1], such as 5-hydroxy-L-tryptophan [2,3], violacein [4,5], indigos [6,7], and halogenated tryptophan [8,9], are extensively utilized in the pharmaceutical, cosmetic, food, and animal feed industries. Currently, the primary method for L-tryptophan production is microbial fermentation. In particular, metabolic engineering of *Escherichia coli* for the production of L-tryptophan has been intensively studied over the past few years, establishing *E. coli* as the predominant microbial chassis for industrial L-tryptophan production. Much of these research has focused on optimizing the rate-limiting steps in the L-tryptophan synthesis pathway, enhancing the availability of metabolic precursors such as phosphoenolpyruvate (PEP) and erythrose-4-phosphate (E4P), and blocking competing pathways and the L-tryptophan degradation pathway [10–14]. Furthermore, the development of riboswitch-based biosensors has facilitated the identification of new genetic targets to enhance L-tryptophan production [15]. By integrating various strategies, engineered *E. coli* strains have achieved L-tryptophan production titers of 40 to 55 g/L and yields of 0.15 to 0.23 g/g glucose in fed-batch fermentation [12,13,16]. However, compared to other amino acids such as L-lysine and L-threonine, the titers and yields of L-tryptophan achieved thus far remain relatively low, substantially below the theoretical yield of L-tryptophan. This discrepancy results in elevated production costs, thereby constraining the broader industrial utilization of L-tryptophan. Accordingly, the development of more robust and efficient microbial strains for L-tryptophan production is highly desirable.

*Corynebacterium glutamicum* is another important industrial chassis broadly used for the industrial production of amino acids [17,18]. It offers several advantages over *E. coli* for large-scale amino acid production, including enhanced safety, greater robustness to environmental stress, increased resistance to phage contamination, and the capacity to utilize a wide range of substrates [18–20]. Systems metabolic engineering strategies have been employed to engineer *C. glutamicum* microbial cell factories for the high-level production of L-lysine, L-arginine, L-proline, etc [21–26]. Early efforts to establish L-tryptophan production in *C. glutamicum* during the 1990s employed a strategy combining random mutagenesis with a limited number of rational genetic modifications [27]. This approach resulted in a strain achieving a record L-tryptophan titer of 58 g/L in fed-batch fermentation, highlighting the potential of *C. glutamicum* for high-level production of this amino acid. However, the engineered *C. glutamicum* strain exhibited several limitations, including retarded cell growth and low productivity, potentially attributed to the accumulation of deleterious mutations during repeated rounds of mutagenesis. Furthermore, the strain’s reliance on a large plasmid and the requirement for exogenous L-phenylalanine and L-tyrosine supplementation hindered its suitability for industrial-scale applications. Therefore, the development of a genetically stable and well-defined *C. glutamicum* strain capable of hyperproducing L-tryptophan remains a priority.

In recent years, further elucidation of the crystal structures and functional mechanisms of the key enzymes for L-tryptophan synthesis, such as the chorismate mutase, the DAHP synthase [28], and the N-(5’-phosphoribosyl) anthranilate isomerase-indole-3-glycerol-phosphate synthase [29], has paved the way for the construction of L-tryptophan hyperproducing *C. glutamicum* strains. Moreover, development of novel L-tryptophan high-throughput screening platforms might facilitate large-scale target mining based on diversified libraries [30]. However, several challenges remain to be solved. Firstly, while the rate-limiting steps and metabolic regulation of L-tryptophan biosynthesis in *E. coli* have been extensively studied, these aspects remain largely unexplored in *C. glutamicum*. Additionally, other potential metabolic engineering targets, such as L-tryptophan exporters or importers, have not been clearly elucidated in *C. glutamicum*. To address this knowledge gap, this study employed the integration of a genome-scale model (GEM) and omics analysis to identify potential metabolic targets. Subsequently, systems metabolic engineering strategies were implemented to engineer a wild-type *C. glutamicum* strain for enhanced L-tryptophan production (Fig. 1). These strategies involved debottlenecking the rate-limiting steps, balancing precursor supply, introducing novel L-tryptophan exporters, and repressing competing pathways. This comprehensive engineering approach enabled the construction of a well-defined *C. glutamicum* strain capable of producing L-tryptophan with a titer of 50.5 g/L in 48h with a yield of 0.17 g/g glucose in fed-batch fermentation, without using plasmids or supplementing other essential amino acids. This work lays the foundation for developing an economically viable process for industrial-scale L-tryptophan production.

**Fig. 1.**
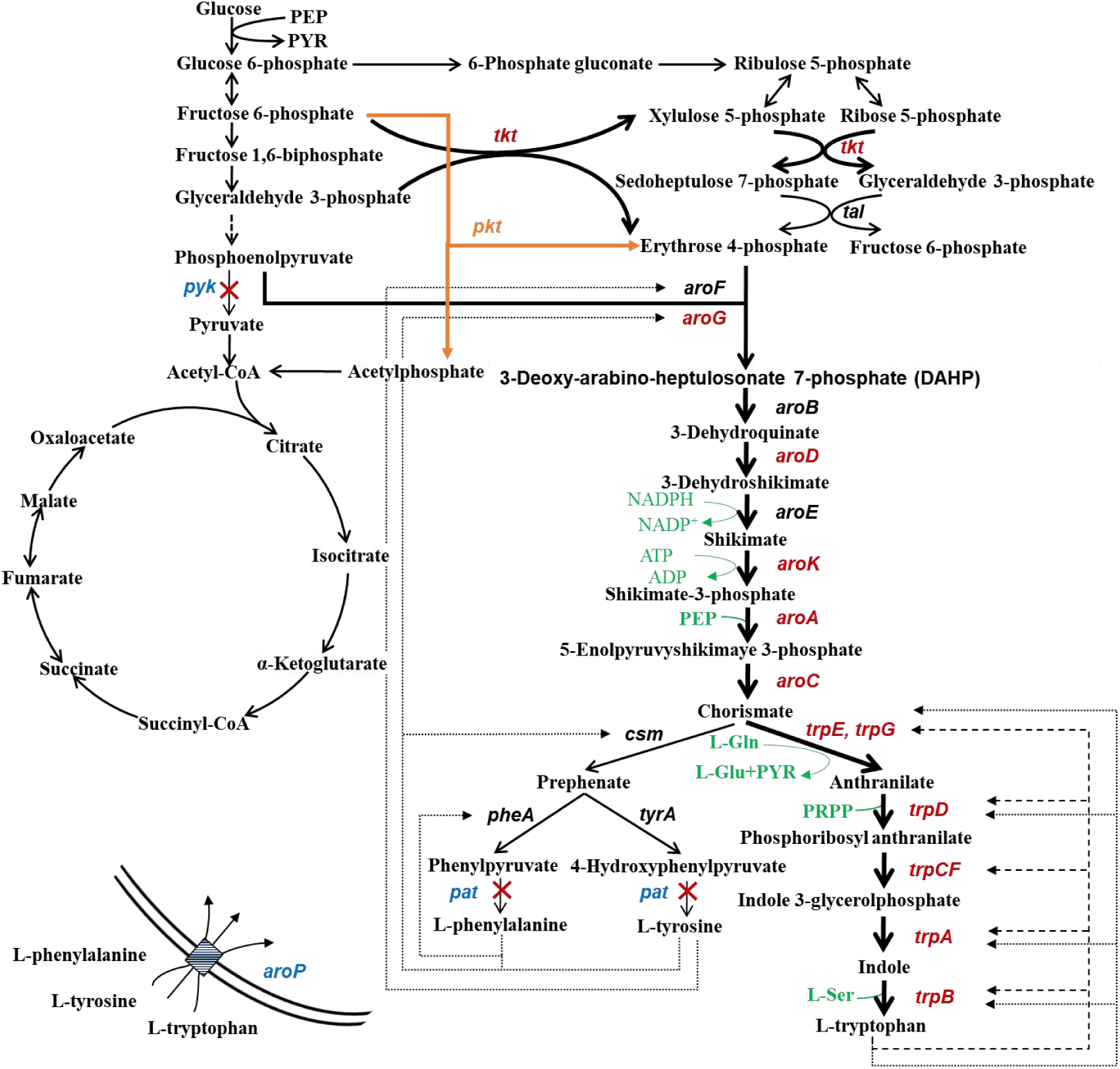
Systematic metabolic engineering of *C. glutamicum* for L-tryptophan production. The dotted lines and dashed lines indicate feedback inhibition and repression, respectively. Precursors and cofactors are colored in green. The overexpression targets are shown with thick lines and red characters. The deletion or down-regulation targets are shown with thin lines and blue characters. The symbol “X” indicates gene deletion. The introduced exogenous phosphoketolase is colored in orange. PEP, phosphoenolpyruvate; PYR, pyruvate; L-Gln, L-glutamine; L-Ser, L-serine; PRPP, phosphoribosyl diphosphate.

## 2. Materials and methods

### 2.1. Bacterial strains and plasmids

The *C. glutamicum* strains used in this study are listed in Table 1. *E. coli* DH5α was used as the host for plasmid construction. The starting *C. glutamicum* strain MB001 was a prophage-free strain derived from ATCC 13032 [31]. The plasmids used in this study are listed in Table S1. The pK18*mobsacB*-derived plasmids were used for markerless gene deletion and integration in *C. glutamicum*. The pEC-K18*mob2* plasmid [32], an *E. coli* and *C. glutamicum* shuttle vector, was applied for gene overexpression in *C. glutamicum* strains.

**Table 1.**
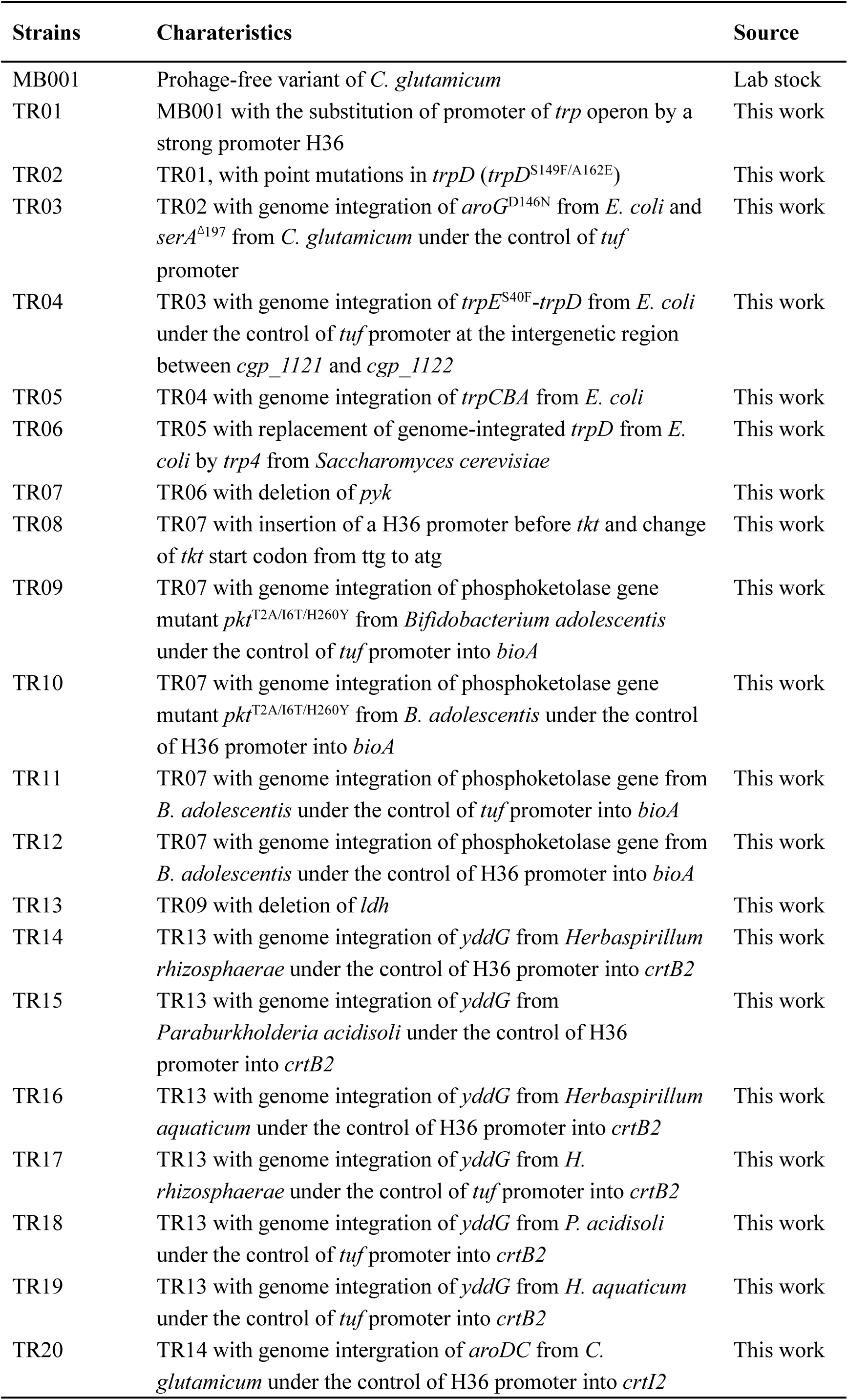
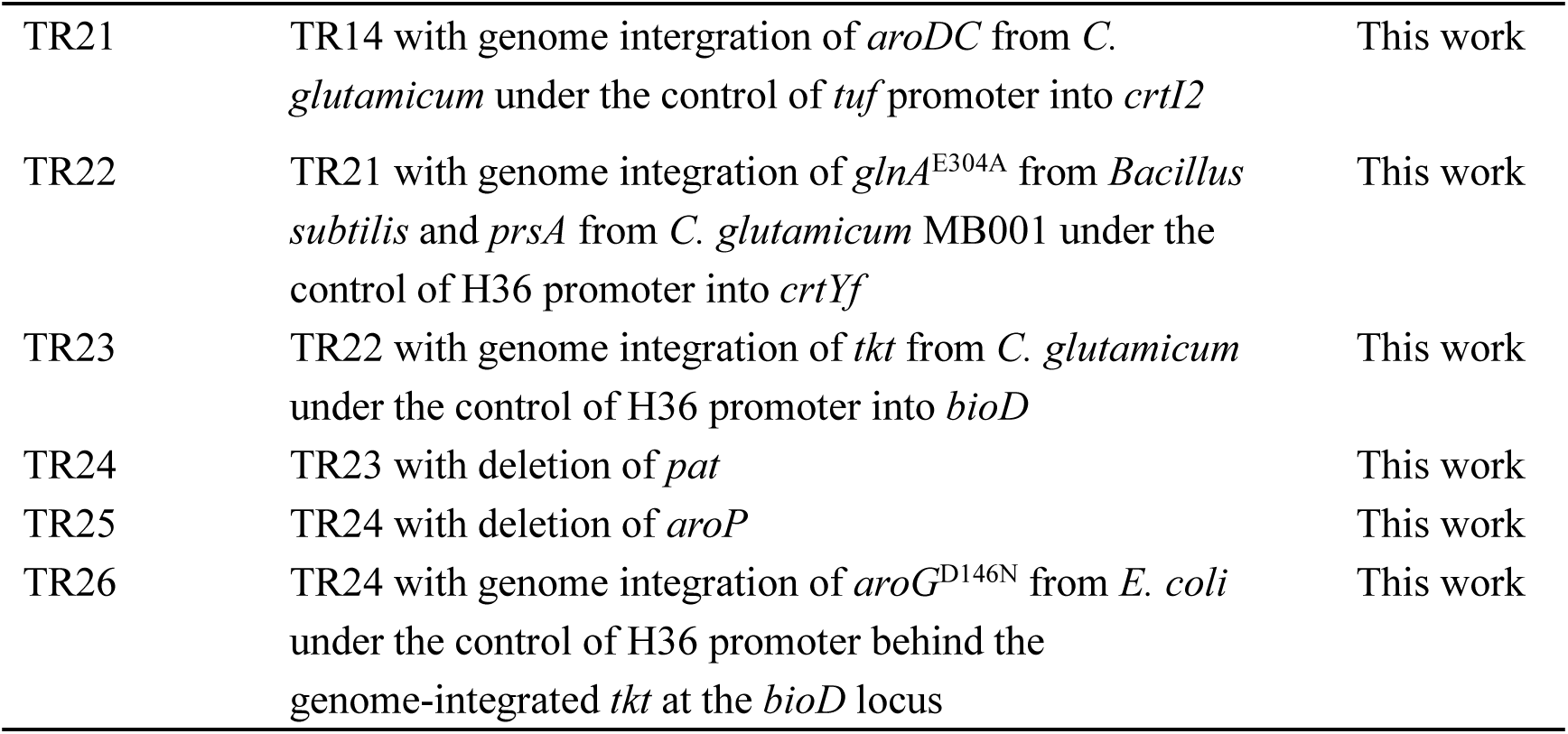
Strains used in this study.

### 2.2. Plasmids and strains construction

Gibson assembly was generally used for plasmid construction following the standard procedures. All of the primers used in this study are listed in Table S2. Gene deletion and insertion in *C. glutamicum* were conducted using the pK18*mobsacB*-derived plasmids through two rounds of recombination [33]. The codon-optimized phosphoketolase gene from *Bifidobacterium adolescentis* and the aromatic amino acid transporter genes were synthesized by Genewiz Company (Suzhou, China). The aromatic amino acid transporters tested in this study are listed in Table S3.

For the construction of plasmids for gene overexpression, the artificial promoter, the functional gene with a common a RBS (aaaggaggttgtc), and the rrnB terminator were inserted into the restriction sites EcoRI/XbaI of the plasmid pEC-K18*mob2* by Gibson assembly.

### 2.3. Genome-scale metabolic modeling

To calculate the optimal metabolic flux distribution for biomass or L-tryptophan biosynthesis, flux balance analysis (FBA) was conducted based on the genome-scale enzyme-constrained model (ecCGL1) [34] of *C. glutamicum* using the CAVE platform [35]. The D-glucose uptake rate was set as 100, and the oxygen uptake rate was set as 1000 for aerobic condition. The objective product was selected as biomass or L-tryptophan.

To predict the specific metabolic engineering targets for L-tryptophan production, protein costs disparities between high growth low product generation (HGLP) (growth rate set as 0.46 h^-1^) and low growth high product generation (LGHP) (growth rate set as 0.1 h^-1^) situations were calculated based on ecCGL1 using ECMpy 2.0 [36]. The glucose uptake rate was set as 10 mmol⋅gDW^-1^ h^-1^. Reactions with fold changes in enzyme costs over 1.5 between LGHP and HGLP conditions were selected as metabolic engineering targets [34]. The calculation was conducted in Python 3.12, and the code was obtained from the previous research [36].

### 2.4. Medium and culture conditions

For shake-flask fermentations, *C. glutamicum* strains were first cultured in 20 mL LSS1 medium in 250 mL baffled shake flasks at 30 ℃, 200 rpm for 14-16 h [37]. The seed cultures were subsequently inoculated into 30 mL LPG2 medium in 500 mL baffled shake flasks with 30 g/L of CaCO_3_ to perform L-tryptophan fermentations [38]. The initial concentration of glucose was 100 g/L. All shake-flask fermentations were conducted in duplicate at 30 ℃, 200 rpm and an initial pH of 7.2. When necessary, 25 μg/L kanamycin was added.

Fed-batch fermentations were carried out in 5 L bioreactors containing 2 L LPG2 at 30 ℃. The pH of the fermentation was kept at 7.0 with the automatic addition of 25% (v/v) ammonia solution. The dissolved oxygen was maintained above 30% of air saturation at an aeration rate of 2 L/min by adjusting the agitation speed. The feeding solution containing 600 g/L glucose was fed to maintain the glucose concentration lower than 10 g/L.

### 2.5. Analysis of cell growth and extracellular metabolites

Cell growth was detected by measuring the optical density of 600 nm (OD_600_) of the samples. The concentrations of glucose, lactic acid, and acetic acid were quantitatively analyzed by high-performance liquid chromatography (HPLC) equipped with an Aminex HPX-87H Column (300×7.8 mm) using 5 mM H_2_SO_4_ as the mobile phase. The flow rate was set as of 0.8 mL/min at 65 ℃ and the injection volume was 20 μL.

The concentrations of L-tryptophan were determined by HPLC with a Dikma Diamonsil AAA Column (5μm, 250×4.6 mm) using the standard phenyl isothiocyanate derivative method.[39] L-tryptophan was separated by using mobile phase A (50 mM CH_3_COONa, pH 6.5) and phase B (methanol: acetonitrile: water=1: 3: 1) with gradient elution at a total flow rate of 1 mL/min at 45 ℃. The detective wavelength was 254 nm and the injection volume was 10 μL.

### 2.6. Metabolome analysis

Intracellular metabolites in *C. glutamicum* cells were extracted as follows. Cells were cultured to the exponential phase and 2 mL of each sample were taken. The cell suspension was centrifuged at 4 ℃, 12000 rpm for 1 min to remove the medium. The cell pellets were washed gently with PBS buffer (pH=7.2-7.4) three times and the supernatant was removed. Afterwards, 1.6 mL of pre-cooled (-80 ℃) 80% methanol was added immediately to resuspend the cell pellets by vortex. The samples were then incubated at -20 ℃ for 30 min and processed by ultra-sonication for 2 min, followed by incubation at -80 ℃ for 2 h for cell lysis. The suspension was subsequently centrifuged at 4 ℃, 14000 rpm for 15 min to remove the protein. The supernatant was taken into a new 2 mL microcentrifuge tube, dried by vacuum and stored at -80 ℃ for further operation.

The metabolites in the supernatant were analyzed by using SCIEXTriple Quad 6500+ liquid chromatography-tandem mass spectrometry (LC-MS/MS) system (AB Sciex, Singapore) equipped with IonDrive detector and Qtrap-6500 mass spectrometer.

## 3. Results and discussion

### 3.1. Identification of potential engineering targets by genome-scale metabolic modeling

As mentioned before, the metabolic regulation and rate-limiting reactions for L-tryptophan biosynthesis in *C. glutamicum* have not been comprehensively elucidated. Genome-scale metabolic modeling is a powerful tool for predicting cellular phenotypes and serves as an invaluable approach for guiding metabolic engineering strategies to enhance chemical production [40]. Recently, the first enzyme-constrained model for *C. glutamicum*, ecCGL1, has been developed [34]. This model incorporates enzyme kinetic data and outperforms the widely used stoichiometry-based model iCW773 in predicting phenotypes and identifying engineering targets [34]. Therefore, we employed ecCGL1 to predict potential engineering targets for L-tryptophan production.

First, the theoretical yield of L-tryptophan and the optimized metabolic flux distribution was calculated based on the stoichiometric data of ecCGL1 using the CAVE platform [35]. As illustrated in Fig. 2A, the theoretical yield of L-tryptophan in *C. glutamicum* is 0.435 mol/mol glucose, which is slightly higher than that in *E. coli* (0.414 mol/mol glucose) [41]. To achieve the theoretical maximum yield of L-tryptophan, the metabolic fluxes toward transketolase reaction involved in E4P biosynthesis, the shikimate pathway, and the terminal steps of L-tryptophan biosynthesis pathway must be significantly enhanced compared to those optimized for cell growth (Fig. 2A). Moreover, the fluxes toward the biosynthesis of key metabolic precursors, including L-serine, L-glutamine, and phosphoribosyl diphosphate (PRPP), should also be increased (Fig. 2A). Conversely, the fluxes involved in the conversion of PEP to pyruvate via the phosphotransferase system (PTS), PEP carboxylase, and pyruvate kinase, as well as flux into tricarboxylic acid (TCA) cycle, must be significantly reduced (Fig. 2A). These predicted results are generally consistent with those previously implemented in *E. coli* [10–14], suggesting that the genome-scale metabolic modeling is reliable.

**Fig. 2.**
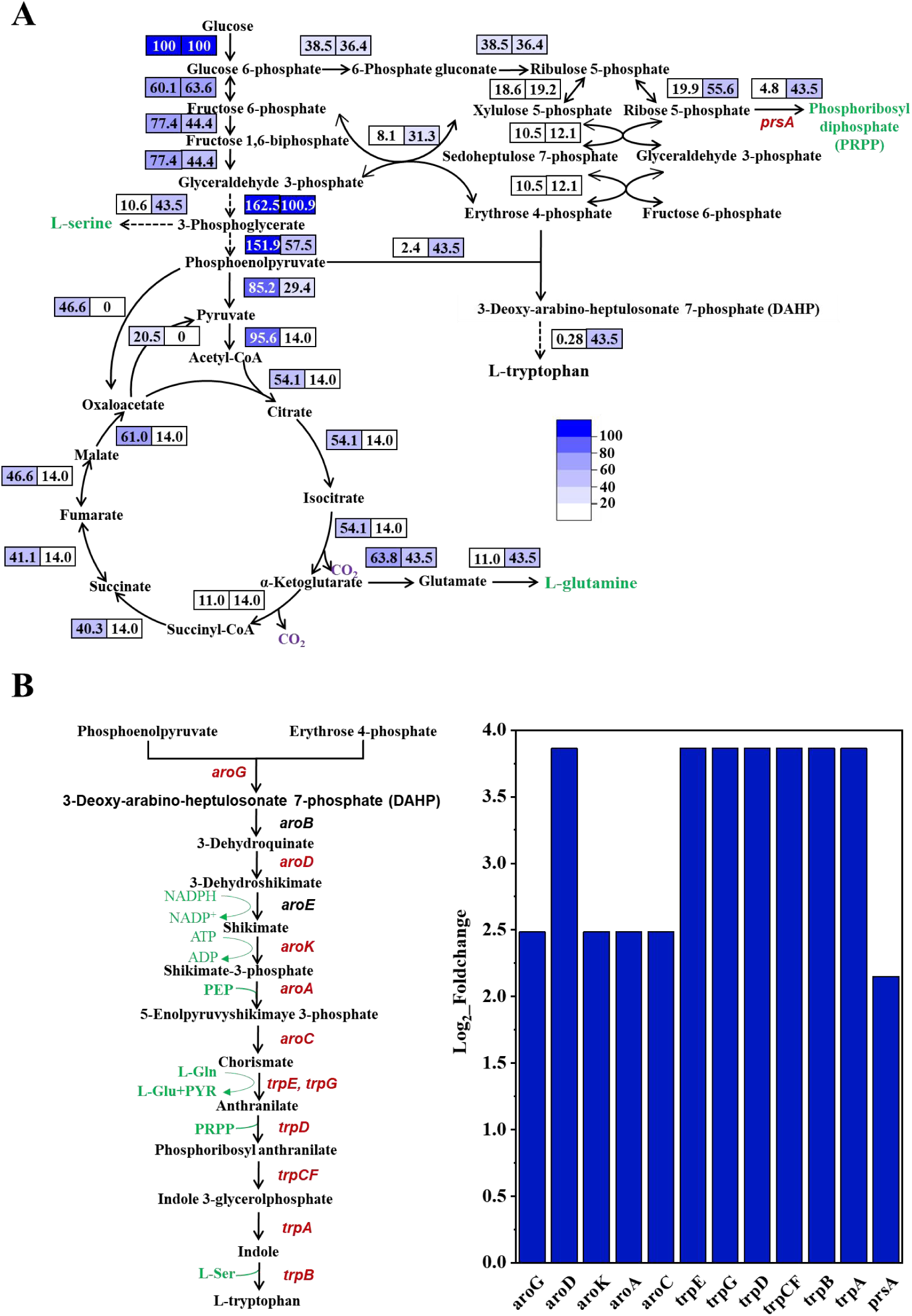
Metabolic engineering targets prediction for L-tryptophan production by using the ecCGL1 model. (A) Metabolic flux distribution with the objective set as biomass accumulation (numbers on the left) and L-tryptophan production (numbers on the right). (B) Enhanced genetic targets identified by the approach of HGLP/LGHP. Precursors and cofactors are colored in green. The predicted targets are colored in red.

While classical FBA based on stoichiometric balance could provide insights into potential up- and down-regulated metabolic pathways, it cannot be used to identify key targets within linear pathways. To overcome this limitation, we employed the ecCGL1 model with the constraints of enzyme kinetic data to pinpoint crucial enzymes based on protein cost discrepancies between HGLP and LGHP conditions [36]. This analysis revealed twelve potential targets with >1.5-fold changes in enzyme cost between HGLP and LGHP states (Fig. 2B and Table S4). These targets could be functionally categorized into three modules: (1) genes (*trpE, trpG*, *trpD*, *trpCF*, *trpB,* and *trpA*) encoding key enzymes within the terminal L-tryptophan biosynthesis pathway, consistent with their roles in L-tryptophan overproduction [16,42]; (2) shikimate pathway genes (*aroG*, *aroD*, *aroK, aroA*, *aroC*), similarly reported to be important for L-tryptophan synthesis in *E. coli* [14]; (3) *prsA* gene encoding ribose-phosphate diphosphokinase, responsible for catalyzing the synthesis of L-tryptophan precursor PRPP. This study further elucidated the specific roles of these genes in modulating L-tryptophan production in *C. glutamicum*.

### 3.2. Enhancement of L-tryptophan synthesis pathway

Genome-scale metabolic modeling indicated that enhanced expression of enzymes within the shikimate and L-tryptophan biosynthetic pathways is crucial for increased L-tryptophan production (Fig. 2B). To enhance the L-tryptophan synthesis pathway, we initially attempted to increase the expression of the entire *trp* operon in *C. glutamicum*. In *E. coli*, the *trp* operon is subject to both feedback repression by the TrpR repressor and transcriptional attenuation mediated by the *trpL* leader sequence (Fig. 3A). While *C. glutamicum* also utilizes *trpL*-mediated attenuation (Fig. 3A), a TrpR homologue is absent in its genome. Therefore, to deregulate the *trp* operon in *C. glutamicum* MB001, the native promoter was replaced with the strong constitutive promoter H36, generating strain TR01. Shake-flask fermentation of TR01 yielded only 0.15 g/L L-tryptophan, demonstrating that increased expression of *trpEGDCFBA* alone is insufficient for substantial overproduction (Fig. 3B). Enzyme-constrained metabolic modeling revealed that anthranilate phosphoribosyl transferase (TrpD) is required with high demand for L-tryptophan biosynthesis (Fig. 2B). Furthermore, unlike its *E. coli* homologue, *C. glutamicum* TrpD is subject to feedback inhibition by L-tryptophan (Fig. 1). To alleviate this inhibition, point mutations S149F/A162E were introduced into TrpD of strain TR01 [43], creating strain TR02. This modification significantly increased L-tryptophan production to 2.70 g/L (Fig. 3B), establishing TrpD as a key control point in *C. glutamicum* L-tryptophan biosynthesis. This contrasts with *E. coli*, where the relief of feedback inhibition by L-tryptophan of anthranilate synthase subunit TrpE rather than TrpD is significant for L-tryptophan production [44].

**Fig. 3.**
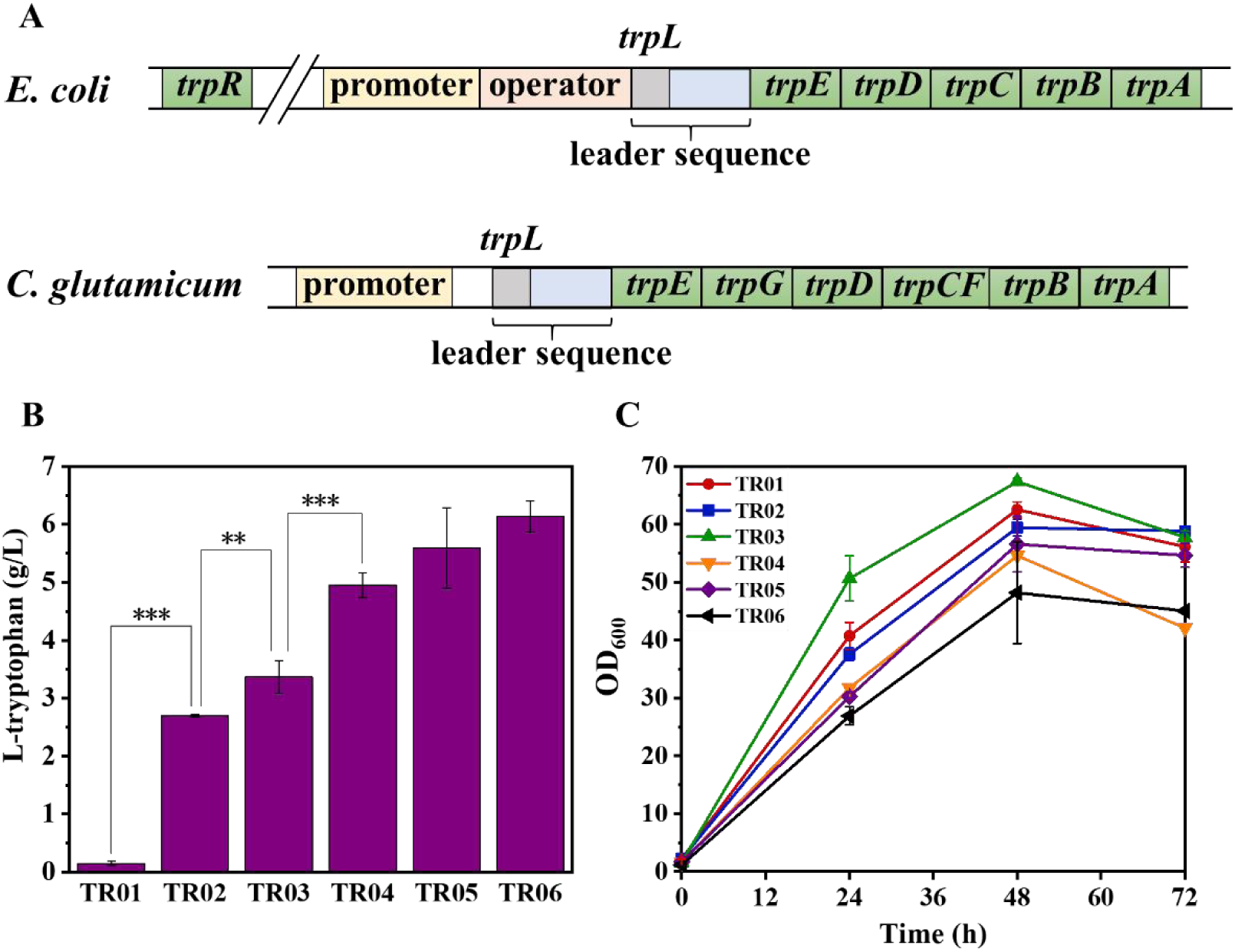
Engineering of L-tryptophan biosynthesis pathway. (A) The *trp* operon in *E. coli* and *C. glutamicum*. (B) L-tryptophan production. (C) Cell growth. Two tailed t-tests indicate statistical significance compared to the parental strain. *P<0.1, **P<0.05, ***P<0.01.

In addition to TrpD, the 3-deoxy-D-arabino-heptulosonate 7-phosphate (DAHP) synthase, which catalyzes the condensation of PEP and E4P to DAHP, is another critical enzyme controlling metabolic flux towards aromatic amino acid biosynthesis (Fig. 2B). Given the feedback inhibition of the native DAHP synthases (AroG and AroF) by aromatic amino acids [45], a feedback-resistant DAHP synthase variant, AroG^D146N^ from *E. coli* [46], was genomically integrated into strain TR02 under the control of tuf promoter. Furthermore, to enhance the biosynthesis of the L-tryptophan precursor L-serine, a copy of feedback-resistant D-3-phosphoglycerate dehydrogenase gene *serA*^Δ197^ from *C. glutamicum* [47] was also integrated after the heterogeneous *aroG*^D146N^ gene. The resulting strain TR03 exhibited an improved L-tryptophan production of 3.37 g/L (Fig. 3B), suggesting that augmenting the supply of precursors via the expression of feedback-resistant *aroG* and *serA* effectively promotes L-tryptophan overproduction [27,48].

To further enhance L-tryptophan production, we sought to optimize the L-tryptophan biosynthesis pathway as informed by enzyme-constrained metabolic modeling (Fig. 2B). Therefore, we attempted to integrated the *trpE*^S40F^-*trpD*-*trpC*-*trpB*-*trpA* gene cluster from *E. coli* into strain TR03. Owing to the considerable size of the gene cluster (>6 kb) and the limited genome integration efficiency in *C. glutamicum*, the operon was divided into two fragments, specifically, *trpE*^S40F^-*trpD* and *trpC*-*trpB*-*trpA*, and sequentially integrated into the genome of TR03 under the control of the *tuf* promoter, resulting in the generation of strains TR04 and TR05, respectively. The *trpE*^S40F^ encodes a feedback-resistant anthranilate synthase. The L-tryptophan production by both TR04 and TR05 was elevated, with strain TR05 achieving a titer of 5.60 g/L (Fig. 3B). Furthermore, given that TrpD was predicted to be a critical target within the L-tryptophan biosynthesis pathway (Fig. 2B), we sought to enhance its activity by incorporating a more efficient variant. Previous studies have reported that the TrpD from *Saccharomyces cerevisiae* encoded by *trp4* gene displays a higher affinity for PRPP and greater enzymatic activity compared to the TrpD from *E. coli* [49]. Consequently, we substituted the *E. coli*-derived *trpD* in TR05 with *trp4* from *S. cerevisiae*, yielding strain TR06. This modification resulted in a further increase in L-tryptophan production to 6.13 g/L (Fig. 3B), confirming its significant role for affecting L-tryptophan overproduction.

### 3.3. Enhancement of PEP and E4P supply

After optimizing the terminal L-tryptophan biosynthesis pathway, efforts were directed towards enhancing the supply of PEP and E4P, the two direct precursors of aromatic amino acids. To increase PEP availability, the *pyk* gene in strain TR06 was deleted, thereby blocking the conversion of PEP to pyruvate via pyruvate kinase. The resulting strain TR07 exhibited a 8.1% increase in L-tryptophan titer (Fig. 4A). Additionally, an increase in both glucose consumption rate (Fig. S1) and L-tryptophan synthesis rate (Fig. 4A) was observed, suggesting that elevating PEP supply positively impacts glucose consumption through the PTS and L-tryptophan biosynthesis via DAHP synthase.

**Fig. 4.**
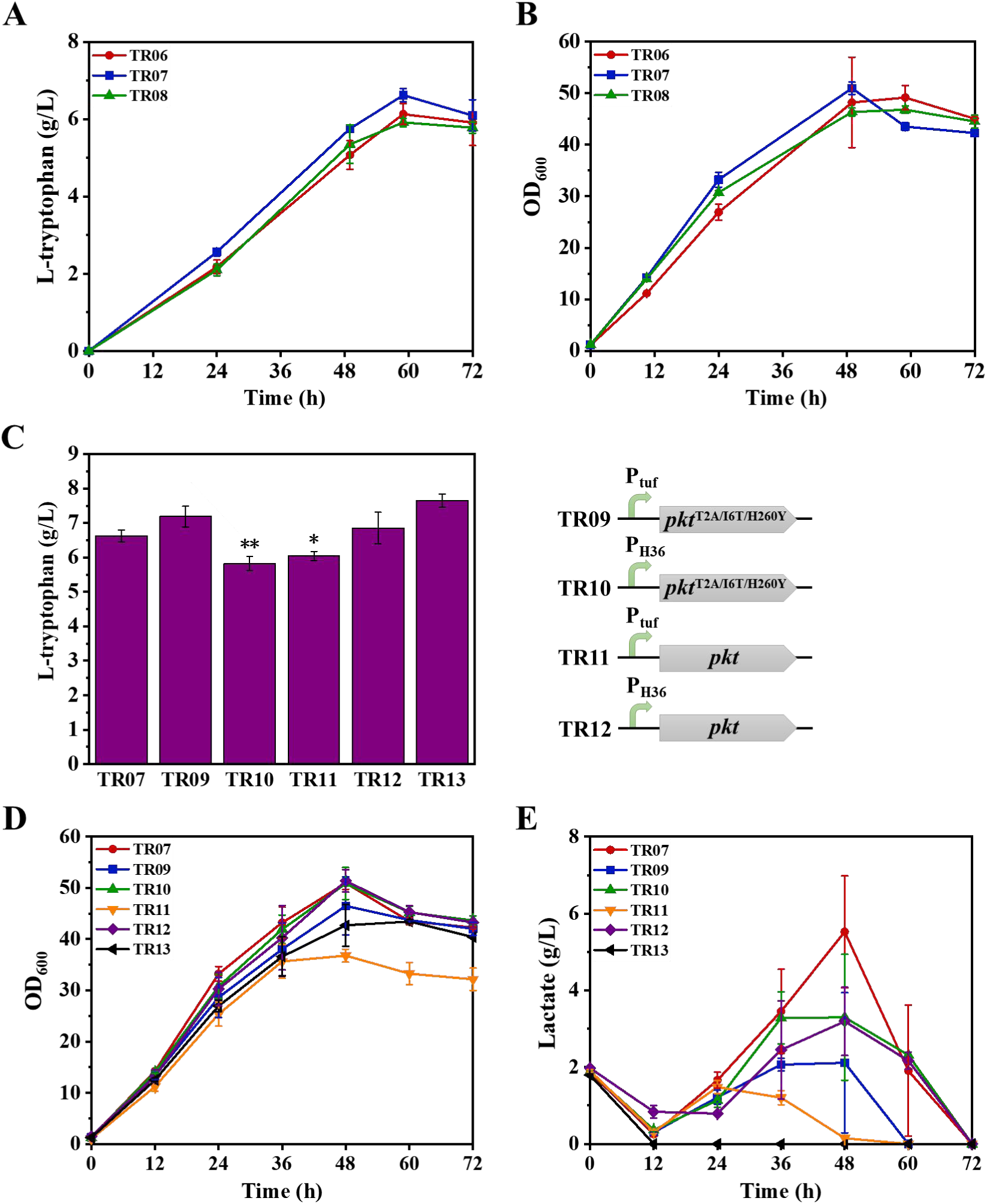
Engineering of the central metabolic pathways. (A) L-tryptophan production by strains TR07 and TR08. (B) Cell growth. (C) L-tryptophan production. (D) Cell growth. (E) Lactate accumulation. Two tailed t-tests indicate statistical significance compared to TR07. *P<0.1, **P<0.05, ***P<0.01.

To further augment E4P supply, two distinct strategies were employed. Initially, we endeavored to boost transketolase activity in strain TR07 by inserting a strong H36 promoter upstream of the endogenous *tkt* gene and substituting its start codon from ttg to atg. However, this modification did not yield an improvement in L-tryptophan yield in the resulting strain TR08 (Fig. 4A). Consequently, an alternative E4P synthesis pathway was explored by introducing a heterologous phosphoketolase. Previous studies have demonstrated that the phosphoketolase from *Bifidobacterium adolescentis* could efficiently convert fructose 6-phosphate (F6P) to E4P and acetylphosphate [50], and this enzyme has been successfully applied to enhance aromatic amino acid production [13]. Consequently, both the wild-type *pkt* gene and its mutant variant *pkt*^T2A/I6T/H260Y^, which encodes an enzyme with higher specific activity [51], were introduced into strain TR07 under the control of either the *tuf* or H36 promoters, generating strains TR09-TR12 (Fig. 4C). These strains exhibited varied responses in terms of cell growth, L-tryptophan production, and lactate accumulation (Fig. 4). Notably, strain TR09 demonstrated the most significant improvement in L-tryptophan titer, showing a 8.4% increase over the control strain TR07 (Fig. 4C). Furthermore, cell growth of TR09 was comparable to that of TR07 (Fig. 4D), while lactate accumulation was notably reduced (Fig. 4E). These findings underscored the importance of optimizing *pkt* expression and phosphoketolase activity to balance cell growth with L-tryptophan production. Given the observed lactate accumulation during L-tryptophan production, the *ldh* gene encoding lactate dehydrogenase was deleted in strain TR09. The resulting strain TR13 exhibited a 6.4% increase in L-tryptophan titer compared to TR09, with no detectable lactate accumulation during fermentation (Fig. 4).

### 3.4. Metabolome analysis

To elucidate alterations in cellular metabolism and potential bottlenecks hindering L-tryptophan production, a comparative metabolome analysis was conducted between the engineered strain TR13 and the wild-type strain MB001. This analysis focused on central metabolic pathway intermediates and intracellular amino acids. As illustrated in Fig. 5A, a total of 108 metabolites within central metabolic pathways were quantified, with significant increases observed in 27 metabolites and significant decreases in 18 metabolites within the TR13 strain (Table S5). Notably, most metabolites in the glycolysis pathway exhibited elevated concentrations in TR13, except for F6P. This exception is possibly due to the introduction of phosphoketolase, which results in the conversion of F6P to E4P. Moreover, the concentrations of PEP and most metabolites in the pentose phosphate pathway were elevated, indicating no apparent supply limitation of PEP and E4P (Fig. S2). Importantly, the concentration of shikimate in TR13 increased by 156-fold, indicating that optimizing the downstream portion of the shikimate pathway to reduce shikimate accumulation is essential for improving L-tryptophan production.

**Fig. 5.**
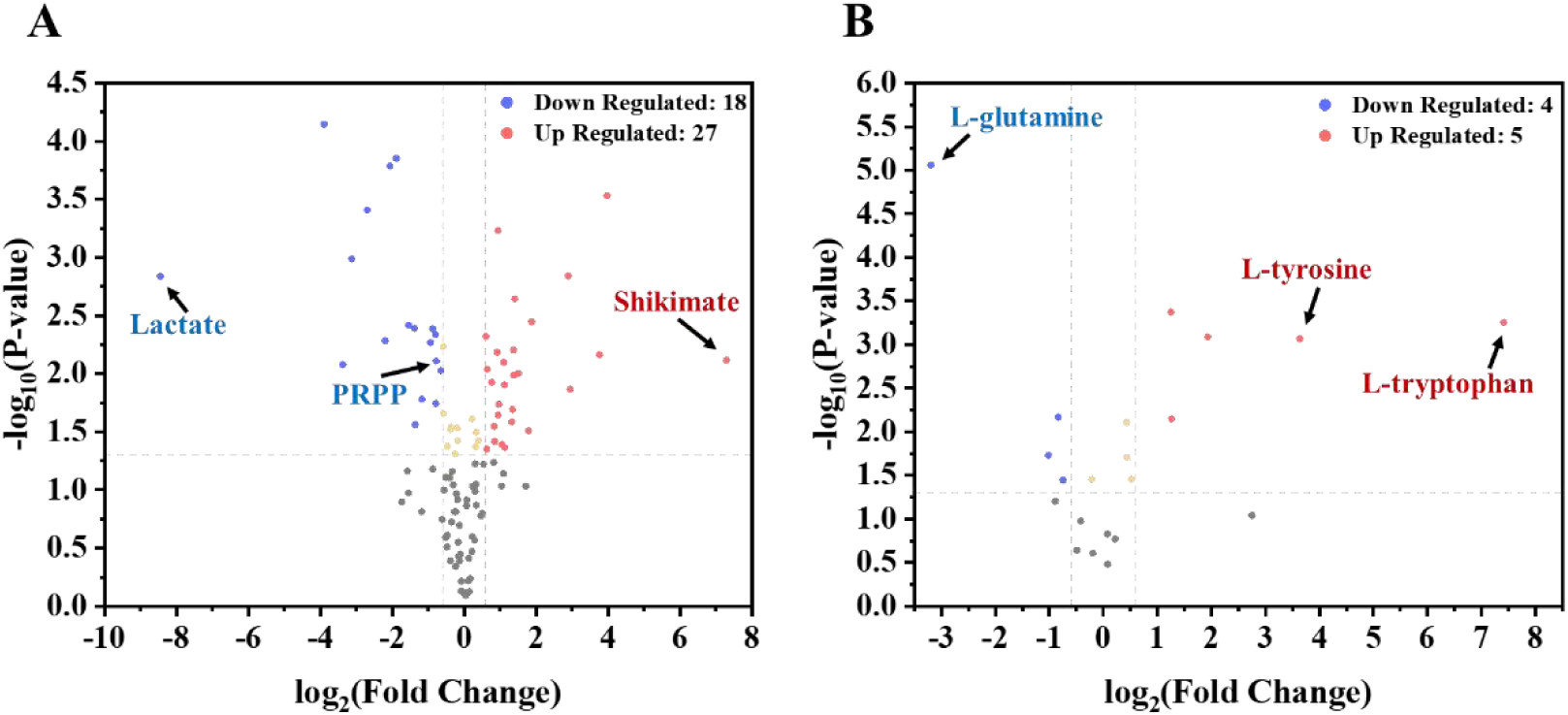
Comparative metabolome analysis of TR13 and MB001. (A) Volcano plot depicting changes of intracellular metabolites within central metabolic pathways. (B) Volcano plot depicting changes of intracellular amino acids. Metabolites with average fold change ≥1.5, t-test p ≤0.05 were considered significantly different.

Regarding the change in intracellular amino acids, TR13 showed significant increases in 5 amino acids and decreases in 4 amino acids compared to MB001 (Fig. 5B and Table S5). In particular, the intracellular levels of L-tryptophan and L-tyrosine in TR13 were elevated by 170-fold and 12.4-fold, respectively. This suggests that enhancing L-tryptophan export and reducing L-tyrosine biosynthesis could be potential strategies for increasing L-tryptophan production. Conversely, a pronounced decrease in the concentrations of L-glutamine and PRPP was observed in TR13, indicating that increasing the availability of these two precursors may be necessary to boost L-tryptophan production. It was also noticed that the concentration of lactate dramatically reduced in TR13 due to deletion of the *ldh* gene. Overall, this metabolome analysis provided new targets for further enhancing L-tryptophan production.

### 3.5. Transporter engineering to increase L-tryptophan production

The metabolome analysis revealed a substantial accumulation of intracellular L-tryptophan in strain TR13. Consequently, enhancing the export of L-tryptophan could mitigate its cellular toxicity and alleviate the feedback inhibition imposed on enzymes in the L-tryptophan biosynthesis pathway. Given that no endogenous L-tryptophan exporter has been identified in *C. glutamicum*, we employed PSI-BLAST to identify potential L-tryptophan exporters in the Uniprot database, using the sequence of the *E. coli* L-tryptophan exporter YddG as a query [15]. Seven candidate exporters were selected (Table S3), and the corresponding genes were subsequently overexpressed in strain TR13 with plasmid pEC-K18*mob2* (Fig. 6A). Strains overexpressing *yddG* from *Herbaspirillum rhizosphaerae* (Hrh), *Herbaspirillum aquaticum* (Haq), and *Paraburkholderia acidisoli* (Pac) exhibited the most significant increase in L-tryptophan titers (Fig. 6A). Subsequently, the three selected *yddG* genes were individually integrated into the genome of strain TR13 under the control of either the H36 or *tuf* promoter, resulting in the generation of strains TR14 to TR19 (Fig. 6C). All engineered strains demonstrated a substantial increase in L-tryptophan titer (Fig. 6C) without compromising cellular growth (Fig. 6B). Specifically, strain TR14, harboring the integrated *H. rhizosphaerae yddG* gene under the H36 promoter, achieved the highest L-tryptophan titer of 11.24 g/L, which is improved by 80.7% compared with strain TR13 (Fig. 6C). These findings suggest that L-tryptophan export represents a critical bottleneck in L-tryptophan production in *C. glutamicum*. Therefore, screening for and optimizing the expression of appropriate *yddG* genes is essential for enhancing L-tryptophan production in this organism.

**Fig. 6.**
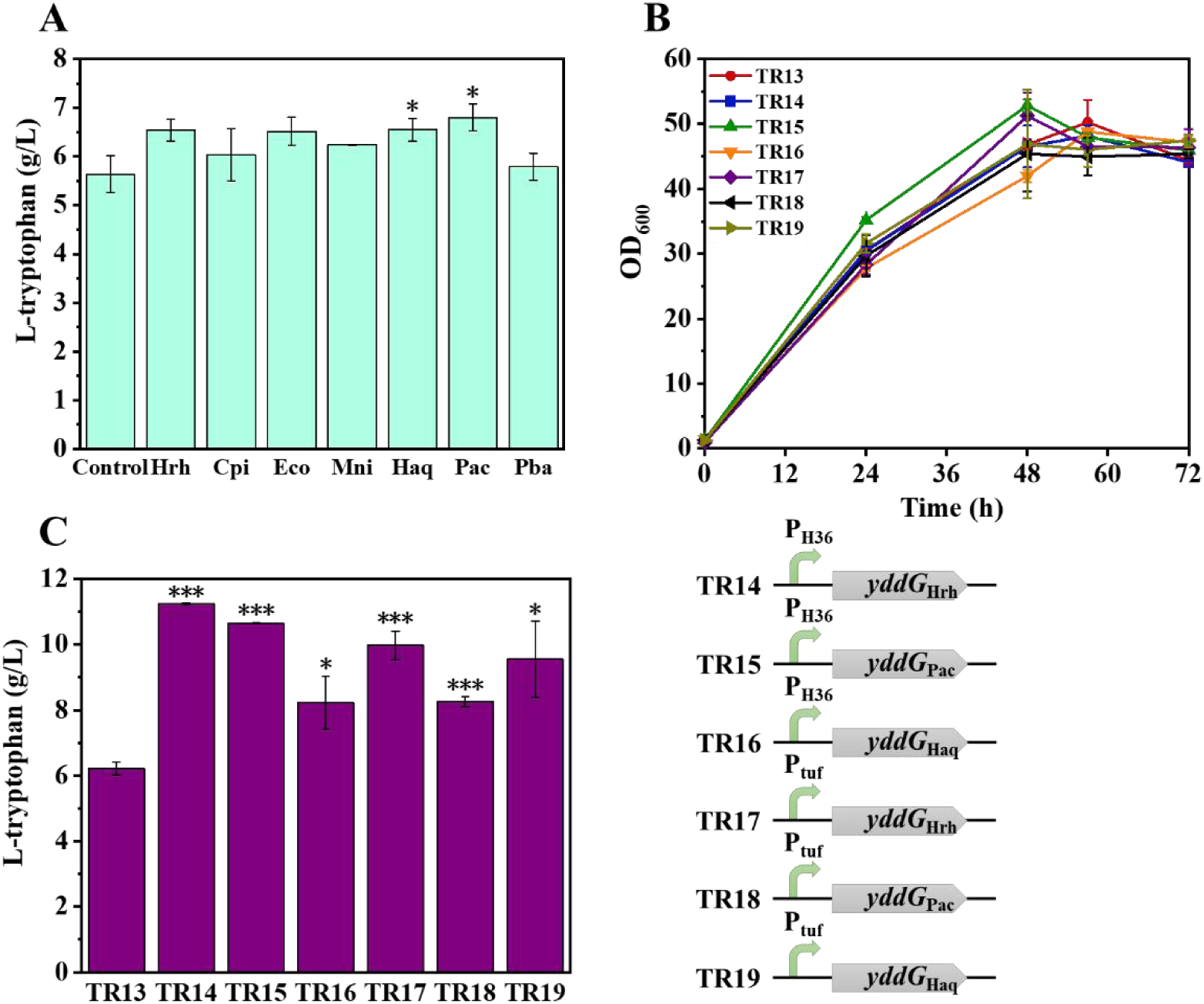
Screening of L-tryptophan exporters. (A) Screening of potential L-tryptophan exporters with plasmid pEC-K18*mob2*. The control group contained an empty pEC-K18*mob2* plasmid. (B) Cell growth of the engineered strains. (C) L-tryptophan production by strains with integrated L-tryptophan exporters in genome. Two tailed t-tests indicate statistical significance compared to the control group or strain TR13. *P<0.1, **P<0.05, ***P<0.01.

### 3.6. Optimization of the shikimate pathway

Given the significant accumulation of shikimate observed in the metabolome analysis of strain TR13 (Fig. 5), it is necessary to optimize the shikimate pathway to redirect metabolic flux toward L-tryptophan production. Enzyme-constrained metabolic modeling indicated that the genes *aroD*, *aroK*, *aroA*, and *aroC* are potential targets within the shikimate pathway (Fig. 2B). To investigate this prediction, each of these genes was individually overexpressed in strain TR13 with plasmid pEC-K18*mob2*. Overexpression of *aroA* did not significantly impact L-tryptophan production, while *aroK* overexpression resulted in a decrease in L-tryptophan titer (Fig. 7A). Conversely, overexpression of either *aroC* or *aroD* led to a 15.8% increase in L-tryptophan titer (Fig. 7A), suggesting that the native expression levels of these genes may limit L-tryptophan biosynthesis in *C. glutamicum*. To further validate the general applicability of these findings, the *aroD* and *aroC* genes were simultaneously integrated into the genome of strain TR14 under the control of either the H36 or tuf promoter, resulting in the creation of strains TR20 and TR21, respectively. Both TR20 and TR21 demonstrated enhanced L-tryptophan production, with strain TR21 achieving the highest titer, a 20.1% improvement compared to the parental strain TR14 (Fig. 7B). Additionally, cell growth of TR20 and TR21 was not negatively affected (Fig. 7C).

**Fig. 7.**
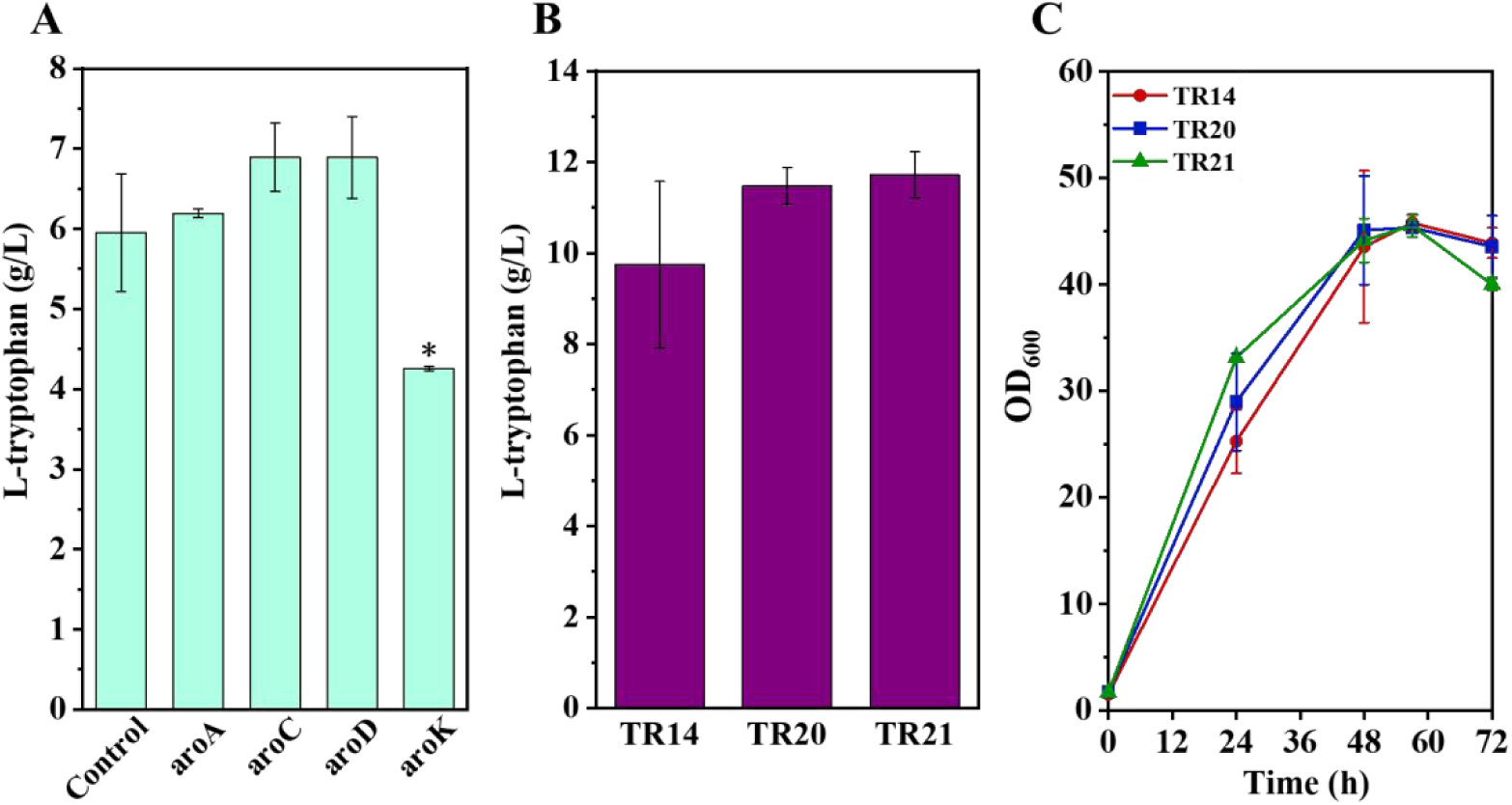
Optimization of the shikimate pathway. (A) L-tryptophan production by strains overexpressing *aroA*, *aroC*, *aroD*, and *aroK* genes. (B) L-tryptophan production by simultaneously overexpressing *aroD*-*aroC* genes. (C) Cell growth of the engineered strains. Two tailed t-tests indicate statistical significance compared to the control group. *P<0.1, **P<0.05, ***P<0.01.

### 3.7. Optimization of the availability of other precursors

Metabolome analysis revealed significantly reduced intracellular concentrations of L-glutamine and PRPP in TR13(Fig. 5), suggesting that increasing the availability of these precursors may enhance L-tryptophan production. To increase the supply of L-glutamine and PRPP, four *glnA* genes encoding glutamine synthetases and three *prsA* genes encoding PRPP synthetase from various organisms (Table S6), which were characterized with high enzymatic activities [52], were individually overexpressed in strain TR14 with plasmid pEC-K18*mob2*. All strains overexpressing *glnA* exhibited increased L-tryptophan production while those overexpressing *prsA* from *E. coli* or *C. glutamicum* also demonstrated elevated L-tryptophan titers (Fig. 8A). The most significant improvements were observed in strains overexpressing the feedback-resistant *glnA*^E304A^ from *Bacillus subtilis* [53] and the native *prsA* from *C. glutamicum* (Fig. 8A). Consequently, the *glnA*^E304A^ gene from *B. subtilis* and the *prsA* gene from *C. glutamicum* were co-integrated into the genome of TR21, creating strain TR22. The L-tryptophan titer by strain TR22 was increased to 12.0 g/L, and the yield was increased from 0.108 to 0.115 g/g glucose, representing a 6.5% improvement compared to strain TR21 (Fig. 8B). Moreover, the cell growth of TR22 was also improved (Fig. 8C).

**Fig. 8.**
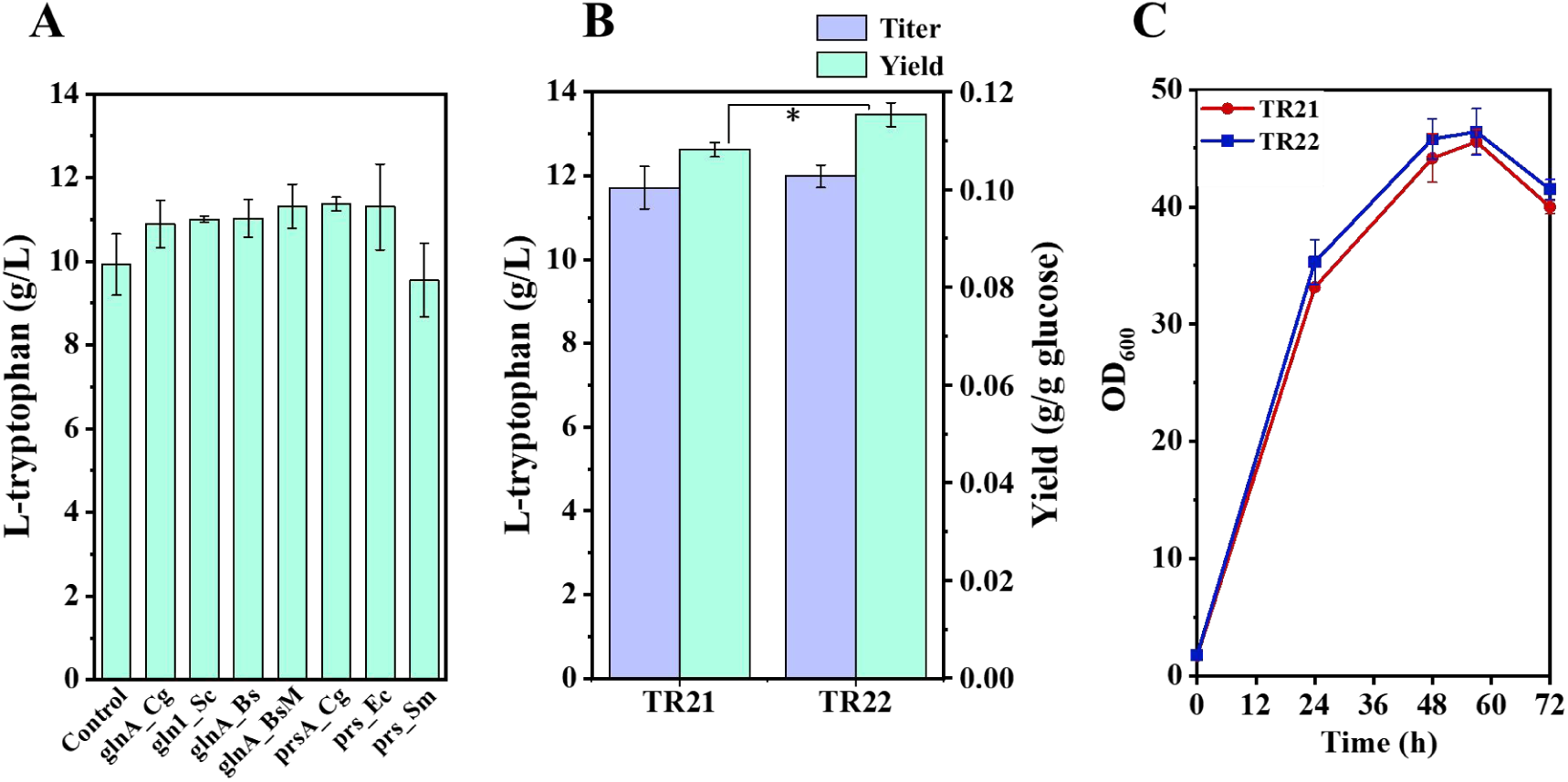
Increasing of the supply of glutamine and PRPP to enhance L-tryptophan production. (A) L-tryptophan production by strains overexpressing glutamine synthetase and PRPP synthetase genes. (B) L-tryptophan production by simultaneously overexpressing *glnA*_BsM-*prsA*_Cg genes. (C) Cell growth of the engineered strains. Two tailed t-tests indicate statistical significance. *P<0.1, **P<0.05, ***P<0.01.

### 3.8. Iterative optimization of key targets and knockdown of competing pathways

Although the production of L-tryptophan by strain TR22 was significantly enhanced through the integration of various metabolic engineering strategies, the resulting titer remains insufficient for industrial application. Recognizing that new bottlenecks may emerge during the engineering process, an iterative optimization approach was employed to fine-tune the supply of precursors and the L-tryptophan biosynthesis pathway. First, to ensure an adequate supply of E4P, an additional copy of the *tkt* gene under the control of the H36 promoter was integrated into the genome of TR22, generating strain TR23. The production of L-tryptophan by strain TR23 increased to 13.7 g/L (Fig. 9A), indicating that E4P shortage had emerged during the construction of strain TR22.

**Fig. 9.**
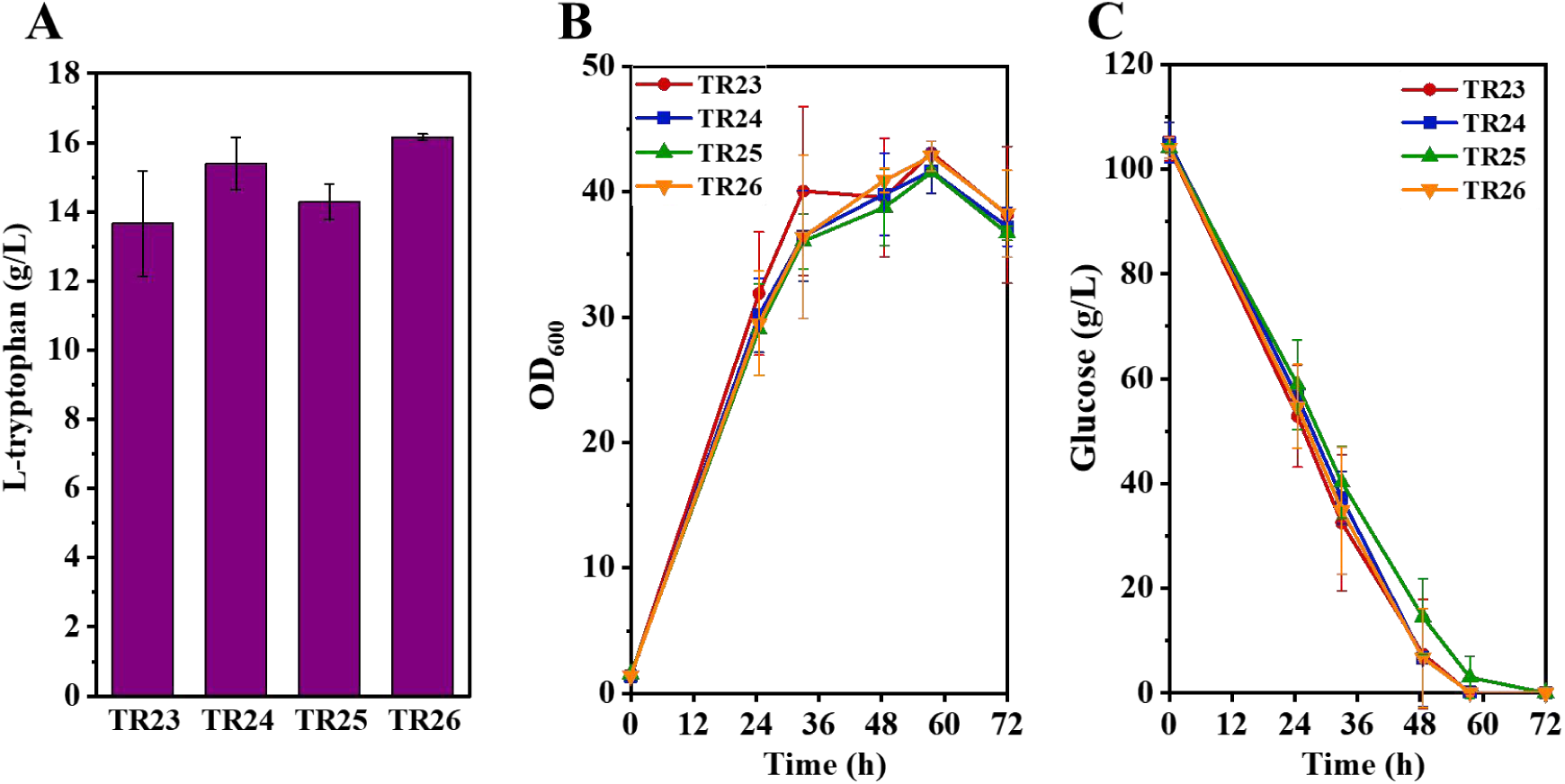
Iterative optimization of the L-tryptophan-producing strain. (A) L-tryptophan production of the engineered strains. (B) Cell growth of the engineered strains. (C) Glucose uptake of the engineered strains.

Subsequently, to reduce the metabolic flux toward the biosynthesis of other aromatic amino acids, the *pat* gene encoding prephenate aminotransferase was deleted in TR23. The resulting strain TR24 exhibited a further increase in L-tryptophan titer to 15.4 g/L (Fig. 9A). Notably, the cell growth of strain TR24 was unaffected (Fig. 9B), which could be due to the existence of other non-specific aminotransferases that can compensate for the synthesis of L-phenylalanine and L-tyrosine [54–56]. To block L-tryptophan uptake, the aromatic amino acid importer gene *aroP* [57] was deleted in strain TR24, generating strain TR25. However, this modification resulted in a slight reduction in L-tryptophan titer (Fig. 9A) as well as decreased glucose consumption rate (Fig. 9C), indicating the import of L-tryptophan is not a limiting factor for L-tryptophan production in *C. glutamicum*.

In an effort to further increase flux through the shikimate pathway, an additional copy of *aroG*^D146N^ from *E. coli* was integrated into the genome of strain TR24. The resulting strain TR26 showed an improved L-tryptophan titer of 16.2 g/L (Fig. 9A) with a yield of 0.16 g/g glucose, which is comparable to the reported yield in *E. coli*. These findings underscore the necessity of iterative optimization in strain engineering to address emerging bottlenecks and achieve higher product titers.

### 3.9. Fed-batch production of L-tryptophan

Fed-batch fermentations of strain TR26 were conducted in 5 L bioreactors to assess its potential for large-scale production of L-tryptophan. L-tryptophan was gradually accumulated during the fermentation process and finally reached a titer of 50.5 g/L at 48 h with a yield of 0.17 g/g glucose (Fig. 10). The obtained titer and yield of L-tryptophan were comparable to the highest levels reported in *E. coli* strains [11–16]. Notably, the biomass of strain TR26 was continuously increased to a high OD_600_ of approximately 280 during the fed-batch fermentation, which might compete with the synthesis of L-tryptophan. Therefore, further genomic modification of strain TR26 to reduce biomass accumulation as well as medium and process optimization could be explored to further increase the titer and yield of L-tryptophan.

**Fig. 10.**
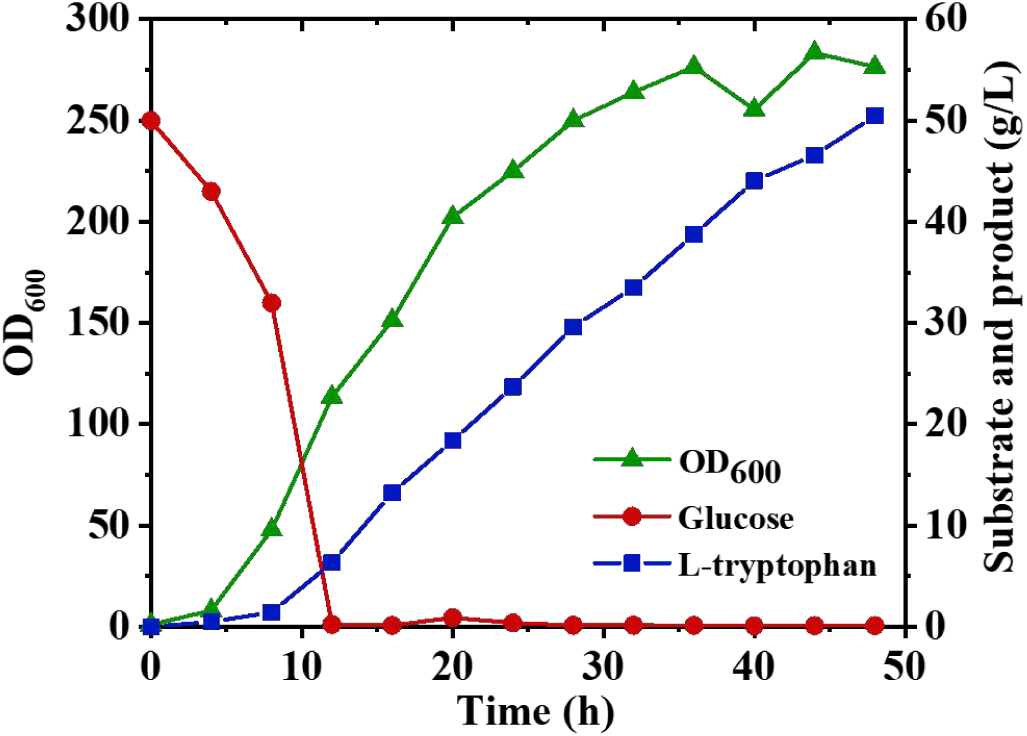
Fed-batch fermentation of strain TR26 in 5 L bioreactor.

## 4. Conclusion

In conclusion, this study successfully engineered L-tryptophan-overproducing *C. glutamicum* strains with a defined genetic background using a systems metabolic engineering approach. By integrating enzyme-constrained genome-scale metabolic modeling (ecCGL1) and comparative metabolome analysis, a series of key targets for enhancing L-tryptophan biosynthesis, including *trpD*, *aroC*, *aroD*, and others, were identified, highlighting differences in metabolic regulation compared to *E. coli*. Furthermore, the overexpression of heterologous phosphoketolase, glutamine synthetases, and PRPP synthetase, coupled with the optimization of selected L-tryptophan exporters, significantly improved precursor supply and export efficiency, leading to a substantial increase in L-tryptophan titer. Through iterative rounds of model-guided engineering and metabolome analysis, a final strain capable of producing 16.2 g/L of L-tryptophan with a yield of 0.16 g/g glucose in shake-flask fermentations was obtained. Fed-batch fermentations of this strain led to an L-tryptophan titer of 50.5 g/L in 48h with a yield of 0.17 g/g glucose, which was comparable to the best reported *E. coli* producers. Notably, the engineered strain is plasmid-free and devoid of unknown mutations, representing a promising platform for industrial L-tryptophan production. Future efforts to further enhance L-tryptophan production can be directed towards the exploration of additional potential targets through multi-omics analysis and the optimization of fermentation processes.

## Acknowledgments

This work was supported by the National Key R&D Program of China (No. 2021YFC2100900), the National Natural Science Foundation of China (Grant Nos. 21938004, 22078172), and Tsinghua University Initiative Scientific Research Program (No. 20223080016).

## Author Contributions

Yufei Dong: Methodology, Investigation, Formal analysis, Data curation, Validation, Writing-original draft preparation. Zhen Chen: Conceptualization, Investigation, Formal analysis, Writing - original draft, Writing - review & editing, Funding acquisition, Resources, Supervision, Project administration.

## Reference

[1] Xiao S, Wang Z, Wang B, Hou B, Cheng J, Bai T, et al. Expanding the application of tryptophan: Industrial biomanufacturing of tryptophan derivatives. Front Microbiol 2023;14:1099098. 10.3389/fmicb.2023.1099098.

[2] Lee J, Lee J. Recent research progress in the microbial production of aromatic compounds derived from L-tryptophan. J Life Sci 2020;30(10):919–29. 10.5352/JLS.2020.30.10.919

[3] Shen YP, Niu FX, Yan ZB, Fong LS, Huang YB, Liu JZ. Recent advances in metabolically engineered microorganisms for the production of aromatic chemicals derived from aromatic amino acids. Front Bioeng Biotechnol 2020;8:407. 10.3389/fbioe.2020.00407.

[4] Sarwar N, Sarwar S, Ejaz S, Al-Adeeb A, Al-Ansi W, Li Y, et al. Metabolic engineering of microorganisms to increase production of violacein. Int J Agric Environ Biotechnol 2021;6(1):295–306. 10.22161/ijeab.61.37.

[5] Füller JJ, Röpke R, Krausze J, Rennhack KE, Daniel NP, Blankenfeldt W, et al. Biosynthesis of violacein, structure and function of l-tryptophan oxidase VioA from *Chromobacterium violaceum*. J Biol Chem 2016;291(38):20068–84. 10.1074/jbc.M116.741561.

[6] Choi KY. A review of recent progress in the synthesis of bio-indigoids and their biologically assisted end-use applications. Dyes Pigm 2020;181:108570. 10.1016/j.dyepig.2020.108570.

[7] Cao M, Gao M, Suástegui M, Mei Y, Shao Z. Building microbial factories for the production of aromatic amino acid pathway derivatives: From commodity chemicals to plant-sourced natural products. Metab Eng 2020;58:94–132. 10.1016/j.ymben.2019.08.008.

[8] Kerbs A, Burgardt A, Veldmann KH, Schäffer T, Lee JH, Wendisch VF. Fermentative production of halogenated tryptophan derivatives with *Corynebacterium glutamicum* overexpressing tryptophanase or decarboxylase genes. Chembiochem 2022;23(9):e202200007. 10.1002/cbic.202200007.

[9] Veldmann KH, Minges H, Sewald N, Lee J-H, Wendisch VF. Metabolic engineering of *Corynebacterium glutamicum* for the fermentative production of halogenated tryptophan. J Biotechnol 2019;291(10):7–16. 10.1016/j.jbiotec.2018.12.008.

[10] Chen L, Zeng AP. Rational design and metabolic analysis of *Escherichiacoli*for effective production of L-tryptophan at high concentration. Appl Microbiol Biotechnol 2016;101(2):559–68. 10.1007/s00253-016-7772-5.

[11] Wang J, Cheng LK, Wang J, Liu Q, Shen T, Chen N. Genetic engineering of *Escherichia coli* to enhance production of l-tryptophan. Appl Microbiol Biotechnol 2013;97(17):7587–96. 10.1007/s00253-013-5026-3.

[12] Guo L, Ding S, Liu Y, Gao C, Hu G, Song W, et al. Enhancing tryptophan production by balancing precursors in *Escherichia coli*. Biotechnol Bioeng 2021;119(3):983–93. 10.1002/bit.28019.

[13] Xiong B, Zhu Y, Tian D, Jiang S, Fan X, Ma Q, et al. Flux redistribution of central carbon metabolism for efficient production of l-tryptophan in *Escherichiacoli*. Biotechnol Bioeng 2021;118(3):1393–404. 10.1002/bit.27665.

[14] Liu S, Wang B-B, Xu JZ, Zhang WG. Engineering of shikimate pathway and terminal branch for efficient production of L-tryptophan in *Escherichia coli*. Int J Mol Sci 2023;24(14):11866. 10.3390/ijms241411866.

[15] Tang M, Pan X, Yang T, You J, Zhu R, Yang T, et al. Multidimensional engineering of *Escherichia coli* for efficient synthesis of L-tryptophan. Bioresour Technol 2023;386:129475. 10.1016/j.biortech.2023.129475.

[16] Niu H, Li R, Liang Q, Qi Q, Li Q, Gu P. Metabolic engineering for improving l-tryptophan production in *Escherichia coli*. J Ind Microbiol Biotechnol 2019;46(1):55–65. 10.1007/s10295-018-2106-5.

[17] Bampidis V, Azimonti G, de Lourdes Bastos M, Christensen H, Dusemund B, Kouba M, et al. Safety and efficacy of l-tryptophan produced by fermentation with *Corynebacterium glutamicum* KCCM 80176 for all animal species. EFSA J 2019;17(6):e05729. 10.2903/j.efsa.2019.5729.

[18] Zha J, Zhao Z, Xiao Z, Eng T, Mukhopadhyay A, Koffas MAG, et al. Biosystem design of *Corynebacterium glutamicum* for bioproduction. Curr Opin Biotechnol 2023;79. 10.1016/j.copbio.2022.102870.

[19] Chai M, Deng C, Chen Q, Lu W, Liu Y, Li J, et al. Synthetic biology toolkits and metabolic engineering applied in *Corynebacteriumglutamicum* for biomanufacturing. ACS Synth Biol 2021;10(12):3237–50. 10.1021/acssynbio.1c00355.

[20] Unthan S, Baumgart M, Radek A, Herbst M, Siebert D, Brühl N, et al. Chassis organism from *Corynebacterium glutamicum*-a top-down approach to identify and delete irrelevant gene clusters. Biotechnol J 2014;10(2):290–301. 10.1002/biot.201400041.

[21] Kogure T, Inui M. Recent advances in metabolic engineering of *Corynebacterium glutamicum*for bioproduction of value-added aromatic chemicals and natural products. Appl Microbiol Biotechnol 2018;102(20):8685–705. 10.1007/s00253-018-9289-6.

[22] Liu J, Xu JZ, Rao ZM, Zhang WG. Industrial production of L-lysine in *Corynebacterium glutamicum*: Progress and prospects. Microbiol Res 2022;262:127101. 10.1016/j.micres.2022.127101.

[23] Zhao Z, Cai M, Liu Y, Hu M, Yang F, Zhu R, et al. Genomics and transcriptomics-guided metabolic engineering *Corynebacterium glutamicum* for l-arginine production. Bioresour Technol 2022;364:128054. 10.1016/j.biortech.2021.125799.

[24] Jiang Y, Sheng Q, Wu XY, Ye BC, Zhang B. L-arginine production in *Corynebacterium glutamicum*: manipulation and optimization of the metabolic process. Crit Rev Biotechnol 2021;41(2):172–85. 10.1080/07388551.2020.1844625.

[25] Zhang J, Qian F, Dong F, Wang Q, Yang J, Jiang Y, et al. De novo engineering of *Corynebacterium glutamicum* for l-Proline production. ACS Synth Biol 2020;9(7):1897–906. 10.1021/acssynbio.0c00249.

[26] Liu J, Liu M, Shi T, Sun G, Gao N, Zhao X, et al. CRISPR-assisted rational flux-tuning and arrayed CRISPRi screening of an l-proline exporter for l-proline hyperproduction. Nat Commun 2022;13(1):891. 10.1038/s41467-022-28501-7.

[27] Ikeda M, Katsumata R. Hyperproduction of tryptophan by *Corynebacteriumglutamicum* with the modified pentose phosphate pathway. Appl Environ Microbiol 1999;65(6):2497–502. 10.1128/AEM.65.6.2497-2502.1999.

[28] Burschowsky D, Thorbjørnsrud HV, Heim JB, Fahrig-Kamarauskaitė Jr, Würth-Roderer K, Kast P, et al. Inter-enzyme allosteric regulation of chorismate mutase in *Corynebacterium glutamicum*: structural basis of feedback activation by trp. Biochemistry 2017;57(5):557–73. 10.1021/acs.biochem.7b01018.

[29] Park W, Son HF, Lee D, Kim IK, Kim KJ. Crystal structure and functional characterization of the bifunctional N-(5’-phosphoribosyl)anthranilate isomerase-indole-3-glycerol-phosphate synthase from *Corynebacteriumglutamicum*. J Agric Food Chem 2021;69(42):12485–93. 10.1021/acs.jafc.1c05132.

[30] Guo H, Wang N, Ding T, Zheng B, Guo L, Huang C, et al. A tRNA Modification-based strategy for Identifying amiNo acid Overproducers (AMINO). Metab Eng 2023;78:11–25. 10.1016/j.ymben.2023.04.012.

[31] Baumgart M, Unthan S, Rückert C, Sivalingam J, Grünberger A, Kalinowski J, et al. Construction of a prophage-free variant of *Corynebacteriumglutamicum* ATCC 13032 for use as a platform strain for basic research and industrial biotechnology. Appl Environ Microbiol 2013;79(19):6006–15. 10.1128/AEM.01634-13.

[32] Tauch A, Kirchner O, Löffler B, Götker S, Pühler A, Kalinowski J. Efficient electrotransformation of *Corynebacteriumdiphtheriae*with a mini-replicon derived from the *Corynebacterium glutamicum* Plasmid pGA1. Curr Microbiol 2002;45(5):362–7. 10.1007/s00284-002-3728-3.

[33] Chen Z, Huang J, Wu Y, Wu W, Zhang Y, Liu D. Metabolic engineering of *Corynebacterium glutamicum* for the production of 3-hydroxypropionic acid from glucose and xylose. Metab Eng 2017;39:151–8. 10.1016/j.ymben.2016.11.009.

[34] Niu J, Mao Z, Mao Y, Wu K, Shi Z, Yuan Q, et al. Construction and analysis of an enzyme-constrained metabolic model of *Corynebacterium glutamicum*. Biomolecules 2022;12(10). 10.3390/biom12101499.

[35] Mao Z, Yuan Q, Li H, Zhang Y, Huang Y, Yang C, et al. CAVE: a cloud-based platform for analysis and visualization of metabolic pathways. Nucleic Acids Res 2023;51(W1):W70–W7. 10.1093/nar/gkad360.

[36] Mao Z, Niu J, Zhao J, Huang Y, Wu K, Yun L, et al. ECMpy 2.0: A Python package for automated construction and analysis of enzyme-constrained models. Synth Syst Biotechnol 2024;9(3):494–502. 10.1016/j.synbio.2024.04.005.

[37] Wu W, Zhang Y, Liu D, Chen Z. Efficient mining of natural NADH-utilizing dehydrogenases enables systematic cofactor engineering of lysine synthesis pathway of *Corynebacterium glutamicum*. Metab Eng 2019;52:77–86. 10.1016/j.ymben.2018.11.006.

[38] Li Z, Dong Y, Liu Y, Cen X, Liu D, Chen Z. Systems metabolic engineering of *Corynebacterium glutamicum* for high-level production of 1,3-propanediol from glucose and xylose. Metab Eng 2022;70:79–88. 10.1016/j.ymben.2022.01.006.

[39] Cohen SA, Strydom DJ. Amino acid analysis utilizing phenylisothiocyanate derivatives. Anal Biochem 1988;174(1):1–16. 10.1016/0003-2697(88)90512-X.

[40] Fang X, Lloyd CJ, Palsson BO. Reconstructing organisms in silico: genome-scale models and their emerging applications. Nat Rev Microbiol 2020;18(12):731–43. 10.1038/s41579-020-00440-4.

[41] Varma A, Boesch BW, Palsson BO. Biochemical production capabilities of *Escherichiacoli*. Biotechnol Bioeng 1993;42(1):59–73. 10.1002/bit.260420109.

[42] Ikeda M, Katsumata R. Tryptophan production by transport mutants of *Corynebacterium glutamicum*. Biosci Biotechnol Biochem 2014;59(8):1600–2. 10.1271/bbb.59.1600.

[43] O’Gara JP, Dunican LK. Mutations in the trpD gene of *Corynebacterium glutamicum* confer 5-methyltryptophan resistance by encoding a feedback-resistant anthranilate phosphoribosyltransferase. Appl Environ Microbiol 1995;61(12):4477–9. 10.1128/aem.61.12.4477-4479.1995.

[44] Caligiuri MG, Bauerle R. Identification of amino acid residues involved in feedback regulation of the anthranilate synthase complex from *Salmonelatyphimurium*. Evidence for an amino-terminal regulatory site. J Biol Chem 1991;266(13):8328–35. 10.1016/s0021-9258(18)92979-0.

[45] Ikeda M. Towards bacterial strains overproducing l-tryptophan and other aromatics by metabolic engineering. Appl Microbiol Biotechnol 2006;69(6):615–26. 10.1007/s00253-005-0252-y.

[46] Kikuchi Y, Tsujimoto K, Kurahashi O. Mutational analysis of the feedback sites of phenylalanine-sensitive 3-deoxy-D-arabino-heptulosonate-7-phosphate synthase of *Escherichia coli*. Appl Environ Microbiol 1997;63(2):761–2. 10.1128/aem.63.2.761-762.1997.

[47] Peters-Wendisch P, Netzer R, Eggeling L, Sahm H. 3-Phosphoglycerate dehydrogenase from *Corynebacteriumglutamicum* : the C-terminal domain is not essential for activity but is required for inhibition by L-serine. Appl Microbiol Biotechnol 2002;60(4):437–41. 10.1007/s00253-002-1161-y.

[48] Ikeda M, Nakanishi K, Kino K, Katsumata R. Fermentative production of tryptophan by a stable recombinant strain of *Corynebacterium glutamicum* with a modified serine-biosynthetic pathway. Biosci Biotechnol Biochem 2014;58(4):674–8. 10.1271/bbb.58.674.

[49] Jun KH, Jang JH, Rim GG, Lee GC, Huang R, Pyo HH. An L-tryptophan-producing microorganism and methods to use it for L-tryptophan production. CN 105143440 A. 2015.

[50] Wang J, Zhang R, Zhang J, Gong X, Jiang T, Sun X, et al. Tunable hybrid carbon metabolism coordination for the carbon-efficient biosynthesis of 1,3-butanediol in *Escherichia coli*. Green Chem 2021;23(21):8694–706. 10.1039/d1gc02867g.

[51] Dele-Osibanjo T, Li Q, Zhang X, Guo X, Feng J, Liu J, et al. Growth-coupled evolution of phosphoketolase to improve l-glutamate production by *Corynebacteriumglutamicum*. Appl Microbiol Biotechnol 2019;103(20):8413–25. 10.1007/s00253-019-10043-6.

[52] Lv Q, Hu M, Tian L, Liu F, Wang Q, Xu M, et al. Enhancing l-glutamine production in *Corynebacterium glutamicum* by rational metabolic engineering combined with a two-stage pH control strategy. Bioresour Technol 2021;341:125799. 10.1016/j.biortech.2021.125799.

[53] Wray LV, Fisher SH. Functional roles of the conserved Glu304 loop of*Ba us uis* glutamine synthetase. J Bacteriol 2010;192(19):5018–25. 10.1128/jb.00509-10.

[54] McHardy AC, Tauch A, Rückert C, Pühler A, Kalinowski J. Genome-based analysis of biosynthetic aminotransferase genes of *Corynebacterium glutamicum*. J Biotechnol 2003;104(1):229–40. 10.1016/S0168-1656(03)00161-5.

[55] Marienhagen J, Kennerknecht N, Sahm H, Eggeling L. Functional analysis of all aminotransferase proteins inferred from the genome sequence of *Corynebacterium glutamicum*. J Bacteriol 2005;187(22):7639–46. 10.1128/jb.187.22.7639-7646.2005.

[56] Kurpejović E, Burgardt A, Bastem GM, Junker N, Wendisch VF, Sariyar Akbulut B. Metabolic engineering of *Corynebacteriumglutamicum* for l-tyrosine production from glucose and xylose. J Biotechnol 2023;363:8–16. 10.1016/j.jbiotec.2022.12.005.

[57] Shang X, Zhang Y, Zhang G, Chai X, Deng A, Liang Y, et al. Characterization and molecular mechanism of AroP as an aromatic amino acid and histidine transporter in *Corynebacterium glutamicum*. J Bacteriol 2013;195:5334–42. 10.1128/JB.00971-13.

