## Supplementary Information for "Systems metabolic engineering of *Corynebacterium glutamicum* for efficient L-tryptophan production"

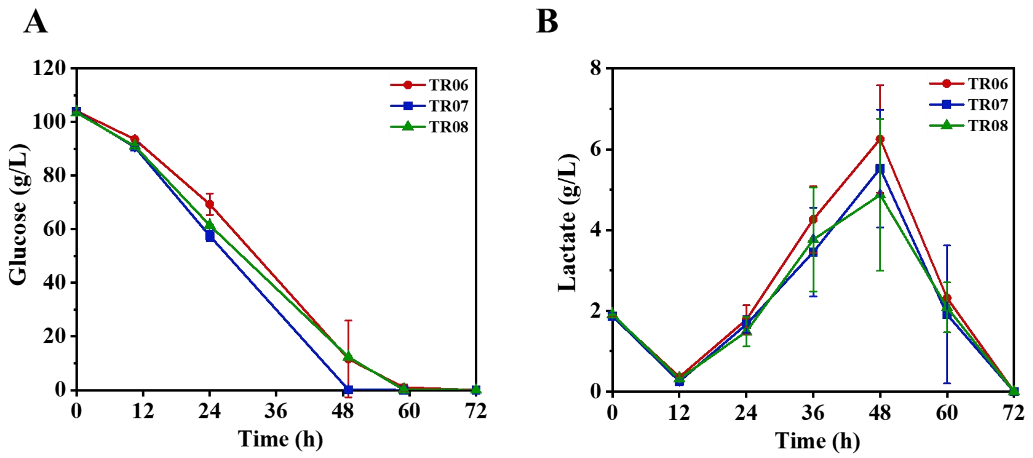


**Fig. S1. Glucose consumption and lactate production profiles of strains with engineered central metabolic pathways.** (A) Glucose consumption. (B) Lactate secretion. All fermentations were conducted in shake flasks in modified LPG2 medium for 72 h.

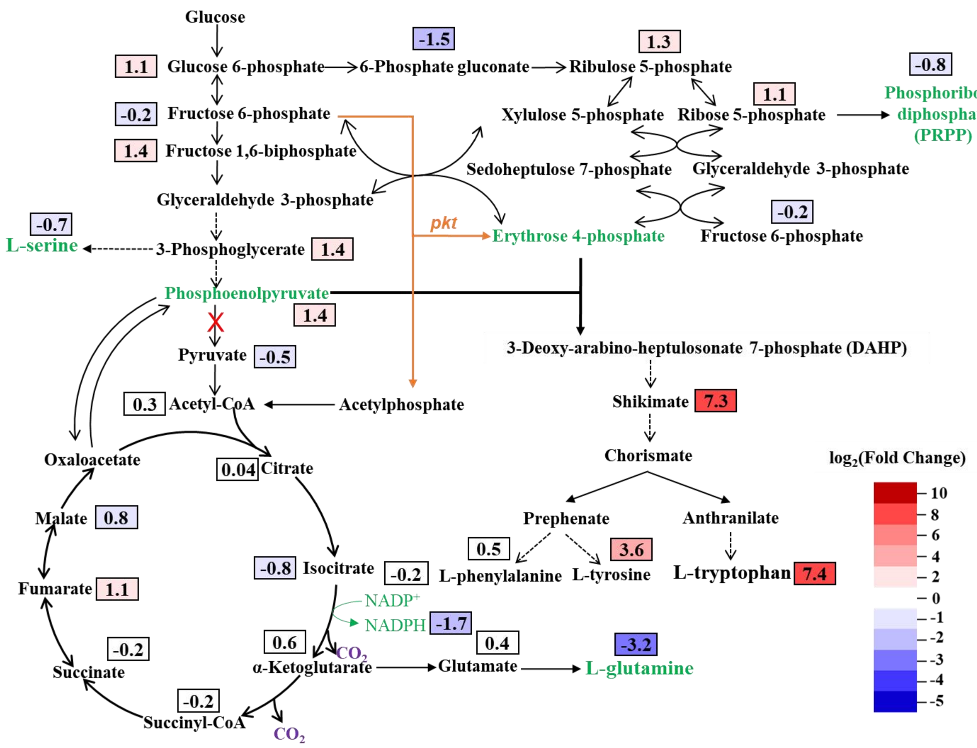


**Fig. S2. Comparative metabolome analysis between *C. glutamicum* strains TR13 and MB001.** The numbers indicate the log2(Fold change) of the intracellular concentrations of the metabolites in TR13 compared with MB001.

**Table S1. Plasmids used in this study**

| Plasmids | Charateristics | Source |
|---|---|---|
| pK18*mobsacB* | Suicide plasmid for gene deletion or integration | Lab stock |
| pEC-K18*mob2* | *E. coli*/*C. glutamicum* shuttle vector | Lab stock |
| pK18-H36-trpE | pK18*mobsacB* derivative for the insertion of an H36 promoter before the *trp* operon | This work |
| pK18-trpDmu | pK18*mobsacB* derivative for integration of *trpD*$^{S149F/A162E}$ | This work |
| pK18-aroG-serA | pK18*mobsacB* derivative for genome integration of *aroG*$^{D146N}$ from *E. coli* and *serA*$^{\Delta 197}$ from *C. glutamicum* | This work |
| pK18-ΔCGP1-tuf-trpED | pK18*mobsacB* derivative for genome integration of *trpE*$^{S40F}$-*trpD* from *E. coli* at the intergenetic region between *cgp_1121* and *cgp_1122* | This work |
| pK18-trpCBA | pK18*mobsacB* derivative for genome integration of *trpCBA* from *E. coli* | This work |
| pK18-ΔtrpD-trp4 | pK18*mobsacB* derivative for genome replacement of *trpD* from *E. coli* by *trp4* from *Saccharomyces cerevisiae* | This work |
| pK18-pykout | pK18*mobsacB* derivative for deletion of *pyk* | This work |
| pK18-H36-ATG-tkt | pK18*mobsacB* derivative for insertion of a H36 promoter before *tkt* and change of *tkt* start codon from ttg to atg | This work |
| pK18-ΔbioA-tuf-PKTWT | pK18*mobsacB* derivative for genome integration of phosphoketolase gene from *Bifidobacterium adolescentis* under the control of *tuf* promoter into *bioA* | This work |
| pK18-ΔbioA-H36-PKTWT | pK18*mobsacB* derivative for genome integration of phosphoketolase gene from *B. adolescentis* under the control of H36 promoter into *bioA* | This work |
| pK18-ΔbioA-tuf-PKTMU | pK18*mobsacB* derivative for genome integration of phosphoketolase gene mutant *pkt*$^{T2A/I6T/H260Y}$ from *B. adolescentis* under the control of *tuf* promoter into *bioA* | This work |
| pK18-ΔbioA-H36-PKTMU | pK18*mobsacB* derivative for genome integration of phosphoketolase gene mutant *pkt*$^{T2A/I6T/H260Y}$ from *B. adolescentis* under the control of H36 promoter into *bioA* | This work |

| | | |
|---|---|---|
| pK18-Δldh | pK18*mobsacB* derivative for deletion of *ldh* | This work |
| pK18-ΔcrtB2-H36-Hrh | pK18*mobsacB* derivative for genome integration of *yddG* from *Herbaspirillum rhizosphaerae* under the control of H36 promoter into *crtB2* | This work |
| pK18-ΔcrtB2-H36-Pac | pK18*mobsacB* derivative for genome integration of *yddG* from *Paraburkholderia acidisoli* under the control of H36 promoter into *crtB2* | This work |
| pK18-ΔcrtB2-H36-Haq | pK18*mobsacB* derivative for genome integration of *yddG* from *Herbaspirillum aquaticum* under the control of H36 promoter into *crtB2* | This work |
| pK18-ΔcrtB2-tuf-Hrh | pK18*mobsacB* derivative for genome integration of *yddG* from *H. rhizosphaerae* under the control of *tuf* promoter into *crtB2* | This work |
| pK18-ΔcrtB2-tuf-Pac | pK18*mobsacB* derivative for genome integration of *yddG* from *P. acidisoli* under the control of *tuf* promoter into *crtB2* | This work |
| pK18-ΔcrtB2-tuf-Haq | pK18*mobsacB* derivative for genome integration of *yddG* from *H. aquaticum* under the control of *tuf* promoter into *crtB2* | This work |
| pK18-ΔcrtI2-tuf-aroDC | pK18*mobsacB* derivative for genome intergration of *aroDC* from *C. glutamicum* under the control of *tuf* promoter into *crtI2* | This work |
| pK18-ΔcrtI2-H36-aroDC | pK18*mobsacB* derivative for genome intergration of *aroDC* from *C. glutamicum* under the control of H36 promoter into *crtI2* | This work |
| pK18-ΔcrtYf-H36-BSglnA(E304A) | pK18*mobsacB* derivative for genome integration of *glnA*$^{E304A}$ from *Bacillus subtilis* into *crtYf* | This work |
| pK18-ΔcrtYf-BSglnA(E304A)-CprsA | pK18*mobsacB* derivative for genome integration of *prsA* from *C. glutamicum* into *crtYf* | This work |
| pK18-ΔbioD-H36-tkt | pK18*mobsacB* derivative for integration of *tkt* from *C. glutamicum* under the control of H36 promoter into *bioD* | This work |
| pK18-Δpat | pK18*mobsacB* derivative for deletion of *pat* | This work |
| pK18-tkt-H36-aroG | pK18*mobsacB* derivative for genome integration of *aroG*$^{D146N}$ from *E. coli* under the control of H36 promoter behind the genome-integrated *tkt* at the *bioD* locus | This work |

| | | |
|---|---|---|
| pK18-ΔaroP | pK18*mobsacB* derivative for deletion of *aroP* | This work |
| pEC-K18-lac-yddG-Hrh | pEC-K18*mob2* containing codon optimized *yddG* from *H. rhizosphaerae* under the control of *lac* promoter | This work |
| pEC-K18-lac-yddG-Cpi | pEC-K18*mob2* containing codon optimized *yddG* from *Cupriavidus pinatubonensis* under the control of *lac* promoter | This work |
| pEC-K18-lac-yddG-Eco | pEC-K18*mob2* containing codon optimized *yddG* from *E. coli* under the control of *lac* promoter | This work |
| pEC-K18-lac-yddG-Mni | pEC-K18*mob2* containing codon optimized *yddG* from *Massilia niastensis* under the control of *lac* promoter | This work |
| pEC-K18-lac-yddG-Haq | pEC-K18*mob2* containing codon optimized *yddG* from *H. aquaticum* under the control of *lac* promoter | This work |
| pEC-K18-lac-yddG-Pac | pEC-K18*mob2* containing codon optimized *yddG* from *P. acidisoli* under the control of *lac* promoter | This work |
| pEC-K18-lac-yddG-Pba | pEC-K18*mob2* containing codon optimized *yddG* from *Proteobacteria bacterium* under the control of *lac* promoter | This work |
| pEC-K18-H36-aroA | pEC-K18*mob2* containing *aroA* from *C. glutamicum* under the control of H36 promoter | This work |
| pEC-K18-H36-aroC | pEC-K18*mob2* containing *aroC* from *C. glutamicum* under the control of H36 promoter | This work |
| pEC-K18-H36-aroD | pEC-K18*mob2* containing *aroD* from *C. glutamicum* under the control of H36 promoter | This work |
| pEC-K18-H36-aroK | pEC-K18*mob2* containing *aroK* from *C. glutamicum* under the control of H36 promoter | This work |
| pEC-K18-lac-CglnA | pEC-K18*mob2* containing *glnA* from *C. glutamicum* under the control of *lac* promoter | This work |
| pEC-K18-lac-gln1 | pEC-K18*mob2* containing *gln1* from *S. cerevisiae* under the control of *lac* promoter | This work |
| pEC-K18-lac-BSglnA | pEC-K18*mob2* containing *glnA* from *B. subtilis* under the control of *lac* promoter | This work |
| pEC-K18-lac-BSglnA(E304A) | pEC-K18*mob2* containing $glnA^{E304A}$ from *B. subtilis* under the control of *lac* promoter | This work |
| pEC-K18-lac-CprsA | pEC-K18*mob2* containing *prsA* from *C.* | This |

| | | |
|---|---|---|
| | *glutamicum* under the control of *lac* promoter | work |
| pEC-K18-lac-Eprs | pEC-K18*mob2* containing *prs* from *E. coli* under the control of *lac* promoter | This work |
| pEC-K18-lac-Sprs | pEC-K18*mob2* containing *prs* from *Serratia marcescens* under the control of *lac* promoter | This work |

**Table S2. Primers used in this study**

| Plasmid | Primer name | Sequence(5'-3') |
|---|---|---|
| pK18-H36-trpE | H1-1.FOR | ataacaatttcacacaggaaacagctatgacatgattacggaagtgacgggtcacaaggacag |
| | H1-1.REV | aggtgcgaccaccctgggcccgttaatattcccccgtttagggcaccagatagaggtacccagcttttgggggcacctaccgaggaaatc |
| | H1-2.FOR | taaacgggggaatattaacgggcccagggtggtcgcaccttggttggtaggagtagcatgggatccatgagcacgaatccccatgttttc |
| | H1-2.REV | acgacggccagtgccaagcttgcatgcctgcaggtcgactcggatatctagctcatcgttgatggag |
| pK18-trpDmu | trpD1.FOR | ataacaatttcacacaggaaacagctatgacatgattacggtcatccgatgacacacgttgt |
| | trpD1.REV | cagggttgtacgcaggtgcgaacaggaaggtgaagttgaacgcttcgaaccacttcacag |
| | trpD2.FOR | caacttcaccttcctgttcgcacctgcgtacaaccctgagattgcgcatgtgcagcc |
| | trpD2.REV | acgacggccagtgccaagcttgcatgcctgcaggtcgactggcgacttcctcctcatcaataacc |
| pK18-aroG-serA | as1.FOR | ataacaatttcacacaggaaacagctatgacatgattacgggatgcctcaacagcctggataac |
| | as1.REV | ctgtgtggatggatctggctgtgagttggacagttgatatccctaaagaag |
| | as2.FOR | atatcaactgtccaactcacagccagatccatccacacagc |
| | as2.REV | atcgtgctatgaataggggttggccgttaccctgcgaatg |
| | as3.FOR | cattcgcagggtaacggccaacccctattcatagcacgatatcgg |
| | as3.REV | acgacggccagtgccaagcttgcatgcctgcaggtcgactggacacaaacctggaatcagcatc |
| pK18-ΔCGP1-tuf-trpED | trpED1.FOR | ataacaatttcacacaggaaacagctatgacatgattacgggatgatgtgatcaccaagactggtatc |
| | trpED1.REV | cattcgcagggtaacggccattcgcgtatggcaatgacagtttgag |
| | trpED2.FOR | ctgtcattgccatacgcgaatggccgttaccctgcgaatgtc |
| | trpED2.REV | gtttaaattgtgtccatgagttaccctcgtgccgccagtg |
| | trpED3.FOR | cactggcggcacgagggtaactcatggacacaatttaaacttcttaaatagcc |
| | trpED3.REV | acgacggccagtgccaagcttgcatgcctgcaggtcgactcgcatcaagcagatctctgagc |
| pK18-trpCBA | trpCBA1.FOR | ataacaatttcacacaggaaacagctatgacatgattacgcagcagaaactagagccagc |
| | trpCBA1.REV | gtttaaattgtgtccatgagttaactgcgcgtcgccgctttc |
| | trpCBA2.FOR | aagcggcgacgcgcagttaactcatggacacaatttaaacttcttaaatagcctc |
| | trpED3.REV | acgacggccagtgccaagcttgcatgcctgcaggtcgactcgcatcaagcagatctctgagc |
| pK18-ΔtrpD-trp4 | trpD41.FOR | ataacaatttcacacaggaaacagctatgacatgattacgctgttctcttat |

| | | |
|---|---|---|
| | | gaccttgtggcgg |
| | trpD41.REV | agcaaagtcgcctcggacatcagaaagtctcctgtgcatgatgc |
| | trpD42.FOR | catgcacaggagactttctgatgtccgaggcgactttgctatc |
| | trpD42.REV | gctaaaacggtttgcatcatctacaaggagctcacactatctataaagtgttc |
| | trpD43.FOR | atagtgtgagctccttgtagatgatgcaaaccgttttagcgaaaatcg |
| | trpD43.REV | acgacggccagtgccaagcttgcatgcctgcaggtcgactttcacgcagcgtatcgatatacagc |
| pK18-pykout | pykout 1.FOR | ataacaatttcacacaggaaacagctatgacatgattacgccatgaccagcgcagattatgc |
| | pykout 1.REV | gacgtactaggcttatgggctcgcttaaatctttcaaaaaatgcgttgacactg |
| | pykout 2.FOR | tttttgaaagatttaagcgagcccataagcctagtacgtcattcc |
| | pykout 2.REV | acgacggccagtgccaagcttgcatgcctgcaggtcgactgtcgcctgactgttaggtttacgttg |
| pK18-H36-ATG-tkt | tkt1.FOR | ataacaatttcacacaggaaacagctatgacatgattacggctgatcaatatcggagtctgccac |
| | tkt1.REV | atagaggtacccagcttttgtccttcctgggttaaaccgggac |
| | tkt2.FOR | ccggtttaacccaggaaggacaaaagctgggtacctctatctggtg |
| | tkt2.REV | ggtgacagcgtcaaggtggtcatggatcccatgctactcctaccaacc |
| | tkt3.FOR | gtaggagtagcatgggatccatgaccaccttgacgctgtcacc |
| | tkt3.REV | acgacggccagtgccaagcttgcatgcctgcaggtcgactggagttcaggcgatcgaacagag |
| pK18-ΔbioA-tuf-PKTWT | Mutuf1.FOR | ataacaatttcacacaggaaacagctatgacatgattacggatgcatccaaggccaatggctag |
| | Mutuf1.REV | cattcgcagggtaacggccagcgcaggtactcggaaatcgc |
| | Mutuf2.FOR | cgatttccgagtacctgcgctggccgttaccctgcgaatgtc |
| | WTtuf2.REV | ccgataaccggagaggtcattgtatgtcctcctggacttcgtgg |
| | WTtuf3.FOR | gaagtccaggaggacatacaatgacctctccggttatcggtaccccgtggaaaaaac |
| | WTtuf5.REV | ccgcgaaacgacggtggatagacaggtggtcttcgttgtc |
| | WTtuf5.FOR | gacaacgaagaccacctgtctatccaccgtcgtttcgcggaac |
| | Mutuf3.REV | cagtggcaatttcatcgacgtcattcgttgtcacccgcggtc |
| | Mutuf4.FOR | ccgcgggtgacaacgaatgacgtcgatgaaattgccactggtttc |
| | Mutuf4.REV | acgacggccagtgccaagcttgcatgcctgcaggtcgactgtttgctgcctcaacgcttaattcag |
| pK18-ΔbioA-H36-PKTWT | Mutuf1.FOR | ataacaatttcacacaggaaacagctatgacatgattacggatgcatccaaggccaatggctag |
| | MuH361.REV | atagaggtacccagcttttggcgcaggtactcggaaatcgc |
| | MuH362.FOR | cgatttccgagtacctgcgccaaaagctgggtacctctatctggtg |
| | WTH362.REV | aaccggagaggtcatggatcccatgctactcctaccaacc |
| | WTH363.FOR | ggttggtaggagtagcatgggatccatgacctctccggttatcggtacccccgtggaaaaaac |
| | WTtuf5.REV | ccgcgaaacgacggtggatagacaggtggtcttcgttgtc |

| | | |
|---|---|---|
| | WTtuf5.FOR | gacaacgaagaccacctgtctatccaccgtcgtttcgcggaac |
| | Mutuf3.REV | cagtggcaatttcatcgacgtcattcgttgtcacccgcggtc |
| | Mutuf4.FOR | ccgcgggtgacaacgaatgacgtcgatgaaattgccactggtttc |
| | Mutuf4.REV | acgacggccagtgccaagcttgcatgcctgcaggtcgactgtttgctgcctcaacgcttaattcag |
| pK18-ΔbioA-tuf-PKTMU | Mutuf1.FOR | ataacaatttcacacaggaaacagctatgacatgattacggatgcatccaaggccaatggctag |
| | Mutuf1.REV | cattcgcagggtaacggccagcgcaggtactcggaaatcgc |
| | Mutuf2.FOR | cgatttccgagtacctgcgctggccgttaccctgcgaatgtc |
| | Mutuf2.REV | ccgataaccggagaggtcattgtatgtcctcctggacttcgtgg |
| | Mutuf3.FOR | gaagtccaggaggacatacaatgacctctccggttatcggtaccccgtggaaaaaac |
| | Mutuf3.REV | cagtggcaatttcatcgacgtcattcgttgtcacccgcggtc |
| | Mutuf4.FOR | ccgcgggtgacaacgaatgacgtcgatgaaattgccactggtttc |
| | Mutuf4.REV | acgacggccagtgccaagcttgcatgcctgcaggtcgactgtttgctgcctcaacgcttaattcag |
| pK18-ΔbioA-H36-PKTMU | Mutuf1.FOR | ataacaatttcacacaggaaacagctatgacatgattacggatgcatccaaggccaatggctag |
| | MuH361.REV | atagaggtacccagcttttggcgcaggtactcggaaatcgc |
| | MuH362.FOR | cgatttccgagtacctgcgccaaaagctgggtacctctatctggtg |
| | MuH362.REV | aaccggagaggccatggatcccatgctactcctaccaacc |
| | MuH363.FOR | ggttggtaggagtagcatgggatccatggcctctccggttaccggtac |
| | Mutuf3.REV | cagtggcaatttcatcgacgtcattcgttgtcacccgcggtc |
| | Mutuf4.FOR | ccgcgggtgacaacgaatgacgtcgatgaaattgccactggtttc |
| | Mutuf4.REV | acgacggccagtgccaagcttgcatgcctgcaggtcgactgtttgctgcctcaacgcttaattcag |
| pK18-Δldh | ldh1.FOR | cacacaggaaacagctatgacatgattacgccgtgatgatgtcatctttgcgtg |
| | ldh1.REV | ctaggcgccaaagattttcgatcccacttcctgatttccc |
| | ldh2.FOR | gaagtgggatcgaaaatctttggcgcctagttggc |
| | ldh2.REV | gtgccaagcttgcatgcctgcaggtcgactggtcgtagttgtgttccaggagg |
| pK18-ΔcrtB2-H36-Hrh | crtBU_F | aggaaacagctatgacatgattacgcacttattgtgcttctcttgggtgcg |
| | crtBH36_R | accagatagaggtacccagcttttgtcatagctgaggctgcttctggt |
| | crtBH36_F | caaaagctgggtacctctatctggtg |
| | PacH36_R | ggatcccatgctactcctaccaacc |
| | H36Hrh_F | ggttggtaggagtagcatgggatccatgaactcaaagaaggcaaccccttatcg |
| | crtBHrh_R | tagtcgagaccttcatcactcaaaggcagtcggcaaccg |
| | HrhcrtB_F | ctttgagtgatgaaggtctcgactaaaactccac |
| | crtBD_R | aagcttgcatgcctgcaggtcgactgaaaaggttgtgatggtcgagctg |
| pK18-ΔcrtB2-H36-Pac | crtBU_F | aggaaacagctatgacatgattacgcacttattgtgcttctcttgggtgcg |
| | crtBH36_R | accagatagaggtacccagcttttgtcatagctgaggctgcttctggt |
| | crtBH36_F | caaaagctgggtacctctatctggtg |

| | | |
|---|---|---|
| | PacH36_R | ggatcccatgctactcctaccaacc |
| | H36Pac_F | ggttggtaggagtagcatgggatccatgacttcaaaaaaaaaagcgacccttatcg |
| | crtBPac_R | gagaccttcatcactcaagagcccttctccgcttcc |
| | PacrtB_F | agggctcttgagtgatgaaggtctcgactaaaactccac |
| | crtBD_R | aagcttgcatgcctgcaggtcgactgaaaaggttgtgatggtcgagctg |
| pK18-ΔcrtB2-H36-Haq | crtBU_F | aggaaacagctatgacatgattacgcacttattgtgcttctcttgggtgcg |
| | crtBH36_R | accagatagaggtacccagcttttgtcatagctgaggctgcttctggt |
| | crtBH36_F | caaaagctgggtacctctatctggtg |
| | PacH36_R | ggatcccatgctactcctaccaacc |
| | H36Haq_F | ggttggtaggagtagcatgggatccatgaagtcaaagaacgcgaccctc |
| | crtBHaq_R | agaccttcatcacctattgattttgagggggaccctcagg |
| | HaqcrtB_F | caaaatcaataggtgatgaaggtctcgactaaaactccac |
| | crtBD_R | aagcttgcatgcctgcaggtcgactgaaaaggttgtgatggtcgagctg |
| pK18-ΔcrtB2-tuf-Hrh | crtBU_F | aggaaacagctatgacatgattacgcacttattgtgcttctcttgggtgcg |
| | tufcrtB_R | ggtaacggccatcatagctgaggctgcttctggt |
| | crtBtuf_F | agcctcagctatgatggccgttaccctgcgaatgtc |
| | Hrhtuf_R | ttgagttcattgtatgtcctcctggacttcgtgg |
| | tufHrh_F | ccaggaggacatacaatgaactcaaagaaggcaacccttatcg |
| | crtBHrh_R | gaccttcatcactcaaaggcagtcggcaaccg |
| | HrhcrtB_F | cgactgcctttgagtgatgaaggtctcgactaaaactccac |
| | crtBD_R | aagcttgcatgcctgcaggtcgactgaaaaggttgtgatggtcgagctg |
| pK18-ΔcrtB2-tuf-Pac | crtBU_F | aggaaacagctatgacatgattacgcacttattgtgcttctcttgggtgcg |
| | tufcrtB_R | ggtaacggccatcatagctgaggctgcttctggt |
| | crtBtuf_F | agcctcagctatgatggccgttaccctgcgaatgtc |
| | Pactuf_R | ttttttttttgaagtcattgtatgtcctcctggacttcgtgg |
| | tufPac_F | acatacaatgacttcaaaaaaaaaagcgacccttatcg |
| | PacrtB2_R | gagaccttcatcactcaagagcccttctccgcttcc |
| | PacrtB2_F | agggctcttgagtgatgaaggtctcgactaaaactccac |
| | crtB2D_R | aagcttgcatgcctgcaggtcgactgaaaaggttgtgatggtcgagctg |
| pK18-ΔcrtB2-tuf-Haq | crtBU_F | aggaaacagctatgacatgattacgcacttattgtgcttctcttgggtgcg |
| | tufcrtB_R | ggtaacggccatcatagctgaggctgcttctggt |
| | crtBtuf_F | agcctcagctatgatggccgttaccctgcgaatgtc |
| | Haqtuf_R | ttcattgtatgtcctcctggacttcgtgg |
| | tufHaq_F | gaagtccaggaggacatacaatgaagtcaaagaacgcgaccctc |
| | crtBHaq_R | agaccttcatcacctattgattttgagggggaccctcagg |
| | HaqcrtB_F | caaaatcaataggtgatgaaggtctcgactaaaactccac |
| | crtBD_R | aagcttgcatgcctgcaggtcgactgaaaaggttgtgatggtcgagctg |
| pK18-ΔcrtI2-tuf-aroDC | crtI2-U.FOR | aggaaacagctatgacatgattacgcactgggcgtgaagaagcaattg |
| | crtI2tuf-U.REV | agggtaacggccaaccacaattcggtcagtgaggtc |
| | crtI2tuf.FOR | ccgaattgtggttggccgttaccctgcgaatgtc |
| | crtI2tuf.REV | atttttccaggcattgtatgtcctcctggacttcgtgg |
| | tufDC.FOR | gaggacatacaatgcctggaaaaattctcctcctcaac |

| | | |
|---|---|---|
| | crtI2-DC.REV | ccggaatagaagattacgctccatcctctaaaccttcgaatttc |
| | crtI2-D.FOR | gatggagcgtaatcttctattccggtgccaccacc |
| | crtI2-D.REV | aagcttgcatgcctgcaggtcgactccatcatgactacggcttttctggc |
| pK18-ΔcrtI2-H36-aroDC | crtI2-U.FOR | aggaaacagctatgacatgattacgcactgggcgtgaagaagcaattg |
| | crtI2-U.REV | tacccagcttttgaccacaattcggtcagtgagg |
| | crtI2-DC.FOR | ccgaattgtggtcaaaagctgggtacctctatctggtg |
| | crtI2-DC.REV | ccggaatagaagattacgctccatcctctaaaccttcgaatttc |
| | crtI2-D.FOR | gatggagcgtaatcttctattccggtgccaccacc |
| | crtI2-D.REV | aagcttgcatgcctgcaggtcgactccatcatgactacggcttttctggc |
| pK18-ΔcrtYf-H36-BSglnA(E304A) | UYf-F | aggaaacagctatgacatgattacggaagcaatggtcaccctggttctc |
| | H36Yf-R | tacccagcttttgctctaggggggattaaaccgagccaaatg |
| | YfH36-F | atccccctagagcaaaagctgggtacctctatctggtg |
| | glnH36-R | tgtactttgccatggatcccatgctactcctaccaacc |
| | H36gln-F | agcatgggatccatggcaaagtacactagagaagatatcg |
| | Dgln-R | gtagtgatgcttcttaatactgagacatatactgttcgcgttcc |
| | glnD-F | tctcagtattaagaagcatcactactagatccaccaccac |
| | DYf-R | aagcttgcatgcctgcaggtcgactcgtgggttgtggagcttgaacg |
| pK18-ΔcrtYf-BSglnA(E304A)-CprsA | Inprs1-F | aggaaacagctatgacatgattacgacccgcgaaacaacttaaaacgg |
| | Inprs1-R | tgacaacctcctttacttaatactgagacatatactgttcgcgttcc |
| | Inprs2-F | cagtattaagtaaaggaggttgtcatgactgctcactggaaacaaaacc |
| | Yfprs-R | tagtagtgatgcttcttaggcctcgccctcgaagag |
| | prsYf-F | cgaggcctaagaagcatcactactagatccaccaccac |
| | DYf-R | aagcttgcatgcctgcaggtcgactcgtgggttgtggagcttgaacg |
| pK18-ΔbioD-H36-tkt | Dtkt-1.FOR | aggaaacagctatgacatgattacgactggacgcgatgggtatcaaattc |
| | Dtkt-1.REV | tacccagcttttgggtttatttccctttaactgcagcatgaagc |
| | Dtkt-2.FOR | gggaaataaacccaaaagctgggtacctctatctggtg |
| | Dtkt-2.REV | tcaaggtggtcatggatcccatgctactcctaccaacc |
| | Dtkt-3.FOR | agcatgggatccatgaccaccttgacgctgtcac |
| | otkt3.REV | ccggcggtttcttaattaaccgttaatggagtccttggcc |
| | otkt4.FOR | taacggttaattaagaaaccgccggcaaggtg |
| | Dtkt-4.REV | aagcttgcatgcctgcaggtcgactcccgatctagctgaaaccaccatc |
| pK18-Δpat | pat-1.FOR | aggaaacagctatgacatgattacgcttcccgttcacgtcctacttg |
| | pat-1.REV | gactacccagcatgatttacacagtactagcccctatggacac |
| | pat-2.FOR | actgtgtaaatcatgctgggtagtctttggcgttttg |
| | pat-2.REV | aagcttgcatgcctgcaggtcgactgccatctaccagcacgacaatttg |
| pK18-tkt-H36-aroG | TA1-F | aggaaacagctatgacatgattacgcccaacatgggatgcagatgagaag |
| | TA1-R | accagatagaggtacccagcttttgttaaccgttaatggagtccttggcc |
| | H36-F | caaaagctgggtacctctatctggtg |
| | TA2-R | cggcggtttcttaattacccgcgacgcgcttttac |
| | TA3-F | gtcgcgggtaattaagaaaccgccggcaaggtg |
| | Dtkt-4.REV | aagcttgcatgcctgcaggtcgactcccgatctagctgaaaccaccatc |
| pK18-ΔaroP | aroP-1.FOR | aggaaacagctatgacatgattacgcttagttaaaacatgatggaagcg |

| | | |
|---|---|---|
| | | gtcg |
| | aroP-1.REV | ttctatcggcctagtatcaaccgtaaacccacatggcac |
| | aroP-2.FOR | tacggttgatactaggccgatagaaattattctggacgtcatg |
| | aroP-2.REV | aagcttgcatgcctgcaggtcgactgatggattgggccgatggatgtg |
| pEC-K18-lac-yddG-Hrh | Hrh1.FOR | acacaggaaacagctatgaccatgattacgaaaggaggttgtcatgaactcaaagaaggcaacccttatcg |
| | Hrh1.REV | aaacagccactcaaaggcagtcggcaaccg |
| | Hrh2.FOR | cggttgccgactgcctttgagtggctgttttggcggatgagagaag |
| | PEC-rrnB_R | agcttgcatgcctgcaggtcgactagagtttgtagaaacgcaaaaaggcc |
| pEC-K18-lac-yddG-Cpi | Cpi1.FOR | acacaggaaacagctatgaccatgattacgaaaggaggttgtcatgcagtccactcggaaggc |
| | Cpi1.REV | aaacagccactcattgcttccatacacgatcacgaac |
| | Cpi2.FOR | atcgtgtatggaagcaatgagtggctgttttggcggatgagagaag |
| | PEC-rrnB_R | agcttgcatgcctgcaggtcgactagagtttgtagaaacgcaaaaaggcc |
| pEC-K18-lac-yddG-Eco | Eco1.FOR | acacaggaaacagctatgaccatgattacgaaaggaggttgtcatgacacgacaaaaagcaacgctc |
| | Eco1.REV | aaacagccacttaaccacgacgtgtcgccag |
| | Eco2.FOR | tggcgacacgtcgtggttaagtggctgttttggcggatgagagaag |
| | PEC-rrnB_R | agcttgcatgcctgcaggtcgactagagtttgtagaaacgcaaaaaggcc |
| pEC-K18-lac-yddG-Mni | Mni1.FOR | acacaggaaacagctatgaccatgattacgaaaggaggttgtcatgaagaacaccaataaggccaccttg |
| | Mni1.REV | aaacagccacctaatcagccacctgcaagacacg |
| | Mni2.FOR | tcttgcaggtggctgattaggtggctgttttggcggatgagagaag |
| | PEC-rrnB_R | agcttgcatgcctgcaggtcgactagagtttgtagaaacgcaaaaaggcc |
| pEC-K18-lac-yddG-Haq | Haq1.FOR | acacaggaaacagctatgaccatgattacgaaaggaggttgtcatgaagtcaaagaacgcgaccctc |
| | Haq1.REV | aaacagccacctattgattttgagggggaccctcagg |
| | Haq2.FOR | gtccccctcaaaatcaataggtggctgttttggcggatgagagaag |
| | PEC-rrnB_R | agcttgcatgcctgcaggtcgactagagtttgtagaaacgcaaaaaggcc |
| pEC-K18-lac-yddG-Pac | Pac1.FOR | acacaggaaacagctatgaccatgattacgaaaggaggttgtcAtgacttcaaaaaaaaaagcgacccttatcg |
| | Pac1.REV | aaacagccactcaagagcccttctccgcttcc |
| | Pac2.FOR | aagcggagaagggctcttgagtggctgttttggcggatgagagaag |
| | PEC-rrnB_R | agcttgcatgcctgcaggtcgactagagtttgtagaaacgcaaaaaggcc |
| pEC-K18-lac-yddG-Pba | Pba1.FOR | acacaggaaacagctatgaccatgattacgaaaggaggttgtcatgcaggccggagacattaaacg |
| | Pba1.REV | aaacagccacttatgactgcgaggaaaggcgg |
| | Pba2.FOR | gcctttcctcgcagtcataagtggctgttttggcggatgagagaag |

| | | |
|---|---|---|
| | PEC-rrnB_R | agcttgcatgcctgcaggtcgactagagtttgtagaaacgcaaaaaggcc |
| pEC-K18-H36-aroA | PEC-H36.FOR | gacacaggaaacagctatgaccatgattacgcaaaagctgggtacctctatctggtg |
| | aroA1.REV | agacgaatcagacatggatcccatgctactcctaccaacc |
| | aroA2.FOR | agtagcatgggatccatgtctgattcgtctatctctttgcccatttg |
| | aroA2.REV | ccgccaaaacagccacctagccaaccatctcctcccaaac |
| | aroA3.FOR | gagatggttggctaggtggctgttttggcggatgagagaag |
| | PEC-rrnB_R | agcttgcatgcctgcaggtcgactagagtttgtagaaacgcaaaaaggcc |
| pEC-K18-H36-aroC | PEC-H36.FOR | gacacaggaaacagctatgaccatgattacgcaaaagctgggtacctctatctggtg |
| | aroC1.REV | gtagtccatcgaagcatggatcccatgctactcctaccaacc |
| | aroC2.FOR | agtagcatgggatccatgcttcgatggactacagcagg |
| | aroC2.REV | ccgccaaaacagccacttacgctccatcctctaaaccttcg |
| | aroC3.FOR | gaggatggagcgtaagtggctgttttggcggatgagagaag |
| | PEC-rrnB_R | agcttgcatgcctgcaggtcgactagagtttgtagaaacgcaaaaaggcc |
| pEC-K18-H36-aroD | PEC-H36.FOR | gacacaggaaacagctatgaccatgattacgcaaaagctgggtacctctatctggtg |
| | aroD1.REV | gaatttttccaggcatggatcccatgctactcctaccaacc |
| | aroD2.FOR | agtagcatgggatccatgcctggaaaaattctcctcctcaac |
| | aroD2.REV | ccgccaaaacagccacctactttttgagatttgccaggatatcgacc |
| | aroD3.FOR | atctcaaaaagtaggtggctgttttggcggatgagagaag |
| | PEC-rrnB_R | agcttgcatgcctgcaggtcgactagagtttgtagaaacgcaaaaaggcc |
| pEC-K18-H36-aroK | PEC-H36.FOR | gacacaggaaacagctatgaccatgattacgcaaaagctgggtacctctatctggtg |
| | aroK1.REV | gaatttgatcattcatggatcccatgctactcctaccaacc |
| | aroK2.FOR | agtagcatgggatccatgaatgatcaaattcacttagatcatcaatcagatg |
| | aroK2.REV | ccgccaaaacagccacttaatcgatttctagatgatgcaacactgc |
| | aroK3.FOR | ctagaaatcgattaagtggctgttttggcggatgagag |
| | PEC-rrnB_R | agcttgcatgcctgcaggtcgactagagtttgtagaaacgcaaaaaggcc |
| pEC-K18-lac-CglnA | lglnA.FOR | acacaggaaacagctatgaccatgattacgaaaggaggttgtcatggcgtttgaaaccccggaag |
| | glnA2.REV | aaacagccacttagcagtcgaagtacaattcgaattcctg |
| | glnA3.FOR | gaattgtacttcgactgctaagtggctgttttggcggatgagagaag |
| | PEC-rrnB_R | agcttgcatgcctgcaggtcgactagagtttgtagaaacgcaaaaaggcc |
| pEC-K18-lac-gln1 | lGln1.FOR | acacaggaaacagctatgaccatgattacgaaaggaggttgtcatggctgaagcaagcatcgaaaag |
| | Gln12.REV | catccgccaaaacagccacttatgaagattctctttcaaattccttcgtcat |

| | | |
|---|---|---|
| | | gtc |
| | Gln13.FOR | gaatcttcataagtggctgttttggcggatgagagaag |
| | PEC-rrnB_R | agcttgcatgcctgcaggtcgactagagtttgtagaaacgcaaaaaggcc |
| pEC-K18-lac-BSglnA | lBSglnA1.FOR | acacaggaaacagctatgaccatgattacgaaaggaggttgtcatggcaaagtacactagagaagatatcg |
| | BSglnA2.REV | ccgccaaaacagccacttaatactgagacatatactgttcgcgttc |
| | BSglnA3.FOR | tatgtctcagtattaagtggctgttttggcggatgagagaag |
| | PEC-rrnB_R | agcttgcatgcctgcaggtcgactagagtttgtagaaacgcaaaaaggcc |
| pEC-K18-lac-BSglnA(E304A) | lBSglnA1.FOR | acacaggaaacagctatgaccatgattacgaaaggaggttgtcatggcaaagtacactagagaagatatcg |
| | BSglnAMU_R | cataacaaggtgctgcatagccaggaacaagacgtttgtaag |
| | BSglnAMU_F | cttgttcctggctatgcagcaccttgttatgtagcatggagc |
| | BSglnA2.REV | ccgccaaaacagccacttaatactgagacatatactgttcgcgttccc |
| | BSglnA3.FOR | tatgtctcagtattaagtggctgttttggcggatgagagaag |
| | PEC-rrnB_R | agcttgcatgcctgcaggtcgactagagtttgtagaaacgcaaaaaggcc |
| pEC-K18-lac-CprsA | lprsA.FOR | acacaggaaacagctatgaccatgattacgaaaggaggttgtcatgactgctcactggaaacaaaacc |
| | prsA2.REV | tccgccaaaacagccacttaggcctcgccctcgaagag |
| | prsA3.FOR | gggcgaggcctaagtggctgttttggcggatgagagaag |
| | PEC-rrnB_R | agcttgcatgcctgcaggtcgactagagtttgtagaaacgcaaaaaggcc |
| pEC-K18-lac-Eprs | lprs.FOR | cacaggaaacagctatgaccatgattacgaaaggaggttgtcatgcctgatatgaagctttttgctggtaac |
| | prs2.REV | atccgccaaaacagccacttagtgttcgaacatggcagagatcg |
| | prs3.FOR | gttcgaacactaagtggctgttttggcggatgagagaag |
| | PEC-rrnB_R | agcttgcatgcctgcaggtcgactagagtttgtagaaacgcaaaaaggcc |
| pEC-K18-lac-Sprs | lSprs1.FOR | acacaggaaacagctatgaccatgattacgaaaggaggttgtcatgccggatatgaaactgtttgcg |
| | lSprs1.REV | catccgccaaaacagccacttaatgttcaaacatcgcgctaatgctttc |
| | lSprs2.FOR | gtttgaacaTtaagtggctgttttggcggatgagagaag |
| | PEC-rrnB_R | agcttgcatgcctgcaggtcgactagagtttgtagaaacgcaaaaaggcc |

**Table S3 Aromatic amino acid transporters tested in this study**

| Organisms | NCBI references | Symbols |
|---|---|---|
| *Herbaspirillum rhizosphaerae* | WP_050478745.1 | Hrh |
| *Paraburkholderia acidisoli* | WP_158954554.1 | Pac |
| *Herbaspirillum aquaticum* | WP_088757482.1 | Haq |
| *Massilia niastensis* | WP_020652949.1 | Mni |
| *Cupriavidus pinatubonensis* | WP_223999738.1 | Cpi |
| *Proteobacteria bacterium* | MBS0348097.1 | Pba |
| *Escherichia coli* | WP_000198205.1 | Eco |

**Table S4 Genetic targets identified by the HGLP/LGHP method**

| reaction | log2_foldchange | gene/enzyme |
|---|---|---|
| DDPA | 2.49 | *aroG* |
| DHQTi | 3.86 | *aroD* |
| SHKK | 2.49 | *aroK* |
| PSCVT | 2.49 | *aroA* |
| CHORS | 2.49 | *aroC* |
| ANS | 3.86 | *trpE* |
| ANS | 3.86 | *trpG* |
| ANPRT | 3.86 | *trpD* |
| PRAIi | 3.86 | *trpCF* |
| TRPS1 | 3.86 | *trpB* |
| TRPS1 | 3.86 | *trpA* |
| PRPPS | 2.15 | *prsA* |

**Table S5 Intracellular metabolites with significant difference in concentration between TR13 and MB001**

| Compound Name | Average fold change | Log2_foldchange | t-test P | Regulation |
|---|---|---|---|---|
| **Central metabolic pathways** | | | | |
| Shikimic Acid | 156.41 | 7.2892 | 0.00764 | Up |
| L-Tyrosine | 15.69 | 3.9714 | 0.00029 | |
| Fructose 1-Phosphate | 13.49 | 3.7536 | 0.00689 | |
| L-Aspartic Acid | 7.67 | 2.9399 | 0.01363 | |
| Inosinic Acid(IMP) | 7.41 | 2.8896 | 0.00144 | |
| Guanosine Monophosphate(GMP) | 3.65 | 1.8681 | 0.00358 | |
| Glyceric Acid | 3.44 | 1.7833 | 0.03099 | |
| Deoxycytidine Monophosphate(dCMP) | 2.84 | 1.5041 | 0.00997 | |
| Fructose 1,6-Bisphosphate | 2.64 | 1.4001 | 0.00227 | |
| 3-Phosphoglycerate | 2.59 | 1.3739 | 0.01033 | |
| Phosphoenolpyruvic Acid | 2.58 | 1.3695 | 0.00625 | |
| D-Ribulose-5-Phosphate/Xylulose 5-Phosphate | 2.53 | 1.3408 | 0.02028 | |
| Cyclic Adenosine Monophosphate(cAMP) | 2.50 | 1.3206 | 0.02597 | |
| Glucose 6-Phosphate | 2.19 | 1.1291 | 0.04328 | |
| Fumaric Acid | 2.17 | 1.1183 | 0.01248 | |
| Cytidine Monophosphate(CMP) | 2.13 | 1.0940 | 0.00798 | |
| Deoxyadenosine Monophosphate(dAMP) | 2.07 | 1.0524 | 0.04061 | |
| GDP-Mannose | 1.94 | 0.9569 | 0.01838 | |
| Uridine Monophosphate(UMP) | 1.92 | 0.9409 | 0.02271 | |
| S-Adenosylhomocysteine | 1.92 | 0.9386 | 0.00059 | |
| Glycine | 1.89 | 0.9191 | 0.00654 | |
| Malic Acid | 1.80 | 0.8492 | 0.03838 | |
| Galactitol/L-Iditol/Mannitol/Sorbitol | 1.78 | 0.8324 | 0.02837 | |
| 2-Hydroxy-3-(4-Hydroxyphenyl)Propanoate | 1.70 | 0.7646 | 0.01187 | |
| 5-Thymidylic Acid(dTMP) | 1.55 | 0.6362 | 0.00917 | |
| Glyceric Acid 1,3-Biphosphate | 1.54 | 0.6216 | 0.04446 | |
| a-Ketoglutarate | 1.52 | 0.6006 | 0.00478 | |
| Thymidine Triphosphate(dTTP) | 0.64 | -0.6509 | 0.00942 | Down |
| Phosphoribosyl Pyrophosphate(PRPP) | 0.58 | -0.7842 | 0.00781 | |
| Isocitric Acid | 0.58 | -0.7920 | 0.01803 | |
| L-Proline | 0.57 | -0.7988 | 0.00461 | |
| 5-Aminoimidazole-4-Carboxamide Ribotide(AICAR) | 0.54 | -0.8784 | 0.00411 | |
| FAD | 0.52 | -0.9378 | 0.00540 | |
| ADP-Glucose | 0.44 | -1.1753 | 0.01656 | |

| | | | | |
|---|---|---|---|---|
| D-2-Hydroxyglutarate | 0.39 | -1.3673 | 0.02748 | |
| L-Carnitine | 0.38 | -1.3858 | 0.00407 | |
| 6-Phosphogluconic Acid-1 | 0.34 | -1.5402 | 0.00385 | |
| Uridine Diphosphate-N-Acetylglucosamine | 0.27 | -1.8865 | 0.00014 | |
| Pipecolic Acid | 0.24 | -2.0606 | 0.00016 | |
| Lac-Phe | 0.22 | -2.1983 | 0.00520 | |
| L-Glutamine | 0.15 | -2.6997 | 0.00039 | |
| Adenine-Neg | 0.11 | -3.1279 | 0.00103 | |
| N-Carbomoyl-L-Aspartate | 0.10 | -3.3726 | 0.00834 | |
| Tropate | 0.07 | -3.9008 | 0.00007 | |
| L-Lactic Acid | 0.003 | -8.4465 | 0.00146 | |
| **Amino acids** | | | | |
| L-Glutamine | 0.11 | -3.1927 | 0.0000 | Down |
| L-Alanine | 0.49 | -1.0182 | 0.0187 | |
| L-Proline | 0.56 | -0.8352 | 0.0068 | |
| L-Serine | 0.60 | -0.7486 | 0.0359 | |
| L-Histidine | 2.38 | 1.2540 | 0.0004 | Up |
| L-lysine | 2.39 | 1.2597 | 0.0071 | |
| Ornithine | 3.82 | 1.9340 | 0.0008 | |
| L-Tyrosine | 12.43 | 3.6363 | 0.0009 | |
| L-Tryptophan | 170.35 | 7.4123 | 0.0006 | |

**Table S6 Glutamine synthetases and PRPP synthetases used in this study**

| Enzyme name | Organisms | NCBI references | Gene name |
|---|---|---|---|
| Glutamine synthetase | *Corynebacterium glutamicum* | AGT05951.1 | glnA_Cg |
| | *Bacillus subtilis* | NC_000964.3 | glnA_Bs |
| | *Bacillus subtilis* | NC_000964.3 | glnA_BsM ($glnA^{E304A}$) |
| | *Saccharomyces cerevisiae* | NP_015360.2 | gln1_Sc |
| PRPP synthetase | *Corynebacterium glutamicum* | AGT04939.1 | prsA_Cg |
| | *Escherichia coli* | NP_415725.1 | prs_Ec |
| | *Serratia marcescens* | WP_004941061.1 | prs_Sm |
